# Adipose tissue-derived neurotrophic factor 3 regulates sympathetic innervation and thermogenesis in adipose tissue

**DOI:** 10.1101/2020.06.06.137273

**Authors:** Xin Cui, Jia Jing, Rui Wu, Qiang Cao, Fenfen Li, Ke Li, Shirong Wang, Liqing Yu, Gary Schwartz, Huidong Shi, Hang Shi, Bingzhong Xue

## Abstract

Activation of brown fat thermogenesis increases energy expenditure and alleviates obesity. Sympathetic nervous system (SNS) is important in brown/beige adipocyte thermogenesis. Here we discover a novel fat-derived “adipokine” neurotrophic factor neurotrophin 3 (NTF3) and its receptor Tropomyosin receptor kinase C (TRKC) as key regulators of SNS growth and innervation in adipose tissue. NTF3 is highly expressed in brown/beige adipocytes, and potently stimulates sympathetic neuron neurite growth. NTF3/TRKC regulates a plethora of pathways in neuronal axonal growth and elongation. Adipose tissue sympathetic innervation is significantly increased in mice with adipocyte-specific NTF3 overexpression, but profoundly reduced in mice with TRKC haploinsufficiency (TRKC+/-). Increasing NTF3 via pharmacological or genetic approach promotes beige adipocyte development, enhances cold-induced thermogenesis and protects against diet-induced obesity (DIO); whereas TRKC+/- mice or SNS TRKC deficient mice are cold intolerant and prone to DIO. Thus, NTF3 is an important fat-derived neurotrophic factor regulating SNS innervation, energy metabolism and obesity.

## Introduction

Obesity has become a serious health problem that poses as a major risk factor for the development of a panel of metabolic diseases such as insulin resistance/type 2 diabetes, dyslipidemia, hypertension and cardiovascular diseases ^1^. Obesity is caused by a chronic energy imbalance that results from energy intake over energy expenditure ^1^. Adipose tissue is one of the most important organs that regulate energy homeostasis in the body ^2^. There exist two distinct types of adipose tissues: white adipose tissue (WAT) that is specialized in energy storage and brown adipose tissue (BAT) that is unique for energy dissipation ^2, 3^. BAT has the capacity to dissipate energy through adaptive thermogenesis, a biological process in which energy is burned as heat instead of being trapped in ATP ^2, 4^. The thermogenic activity of BAT largely depends on the unique action of uncoupling protein 1 (UCP1) in the inner mitochondrial membrane, which actively uncouples oxidative phosphorylation from ATP synthesis, thereby profoundly increasing energy expenditure ^2, 5^. However, recent studies also discovered UCP1-independent thermogenesis mediated by SERCA2b-induced calcium cycling and creatine-driven substrate cycling ^6, 7^. Although brown fat thermogenesis was traditionally viewed as a defense mechanism against cold in rodents, recent discovery of metabolically active brown fat in adult humans has implicated BAT thermogenesis as a promising therapeutic target for the treatment of obesity ^8, 9, 10^.

There are two kinds of UCP1-positive brown adipocytes identified in rodents: traditional brown adipocytes residing in anatomically defined area (e.g., interscapular (iBAT)) and beige adipocytes dispersed in white fat depots ^2, 5^. Unlike brown adipocytes that develop prenatally and persist throughout life time, beige adipocytes are mostly induced by cold and β-adrenergic agonists ^2, 5^. Xue et al previously reported the existence of a unique subset of beige adipocytes, so called developmentally-induced beige adipocytes, which are transiently induced in postnatal mice ^11^.

Adipose tissue is innervated by SNS that plays a key role in adipose lipolysis and BAT/beige thermogenesis ^12, 13, 14, 15, 16^. Catecholamines released by sympathetic nerve terminals in response to cold stimulate lipolysis and activate BAT/beige thermogenesis via β-adrenergic receptors ^17, 18^. Although extensive studies have been devoted to the role of neuronal network and diverse neural-endocrine pathways in the regulation of SNS activation ^19, 20^, much less is known about the role of SNS-targeted tissues (e.g., BAT and WAT) in regulating the development and activation of SNS. A recent study from Spiegelman’s group demonstrated that S100, a BAT-derived secretory protein, promotes SNS innervation into adipose tissue ^21^. However, the mechanism underlying the neurotrophic effect of S100 on SNS is not entirely clear.

It is noteworthy that the developmentally-induced beige adipocytes appear transiently, peaking at postnatal day 20 and then disappearing thereafter toward adulthood ^11^. However, the mechanisms mediating the induction and disappearance of the developmental beige adipocytes are not known. Here we found that the disappearance of the developmental beige cells in adult mice is associated with the diminished sympathetic innervation in white adipose tissue. Our data indicate that the physiological pathway(s) required to maintain the vigorous innervation of sympathetic nerves could decline in adult mice, leading to the disappearance of the developmental beige adipocytes. We discovered that the expression of the fat-derived neurotropic factor NTF3 is higher in mouse BAT than WAT; NTF3 expression coincides with the appearance of developmentally-induced beige adipocytes in postnatal mice, and is induced in WAT by cold exposure in adult mice. Using pharmacological and genetic approaches we further determined the role of NTF3 and its receptor neurotrophic receptor 3/Tropomyosin receptor kinase C (TRKC) in the regulation of SNS growth and innervation in adipose tissue and energy metabolism during cold- and diet-induced thermogenesis.

## Results

### Adipose-derived neurotrophic factor NTF3 correlates with UCP1 expression and sympathetic innervation in adipose tissue

Xue et al (a senior author in this paper) previously found that beige adipocytes can be induced in WAT not only during cold exposure in adult animals, but also in new born pups, which peaked at 20 days of age and then gradually disappeared and replaced by mature white adipocytes by the time when mice were 2 months of age ^11^. Indeed, here we found that *Ucp1* mRNA in inguinal WAT (iWAT) was dramatically reduced in 3-month- and 6-month-old mice compared to postnatal day 20 (P20) pups (**Fig 1A**). Similar reduction of UCP1 protein levels in iWAT was observed in 3-month adult mice compared to P20 pups (**Fig 1B**). The reduction of UCP1 expression in adult iWAT was associated with reduced expression of the SNS marker Tyrosine hydroxylase (TH) measured by immunoblotting (**Fig 1B**). A similar reduction of UCP1 and TH protein levels were also observed in epidydimal WAT (eWAT) of 3-month-old adult mice compared to that of P20 pups (**Suppl. Fig 1A**). We also established a whole mount adipose tissue clearing approach (Adipo-Clear) to permit more comprehensive and accurate three-dimensional visualization of sympathetic innervation with immunostaining of TH ^22^. Compared to iWAT from P20 pups, iWAT from 3-month-old mice had significantly diminished SNS nerve innervation (**Fig 1C** left panel). Quantitation of SNS innervation in iWAT using Imaris Image Analysis Software revealed that mean nerve fiber density, mean nerve fiber length and mean nerve branching points were significantly reduced in iWAT of 3-month-old mice compared to that of P20 pups (**Fig 1C** right panel). A similarly diminished sympathetic innervation was also observed in eWAT of 3-month-old adult mice compared to that of P20 pups (**Suppl. Fig 1B**). These data suggest that the reduced SNS innervation may lead to the disappearance of the developmental beige adipocytes in adult mice.

**Figure 1.**
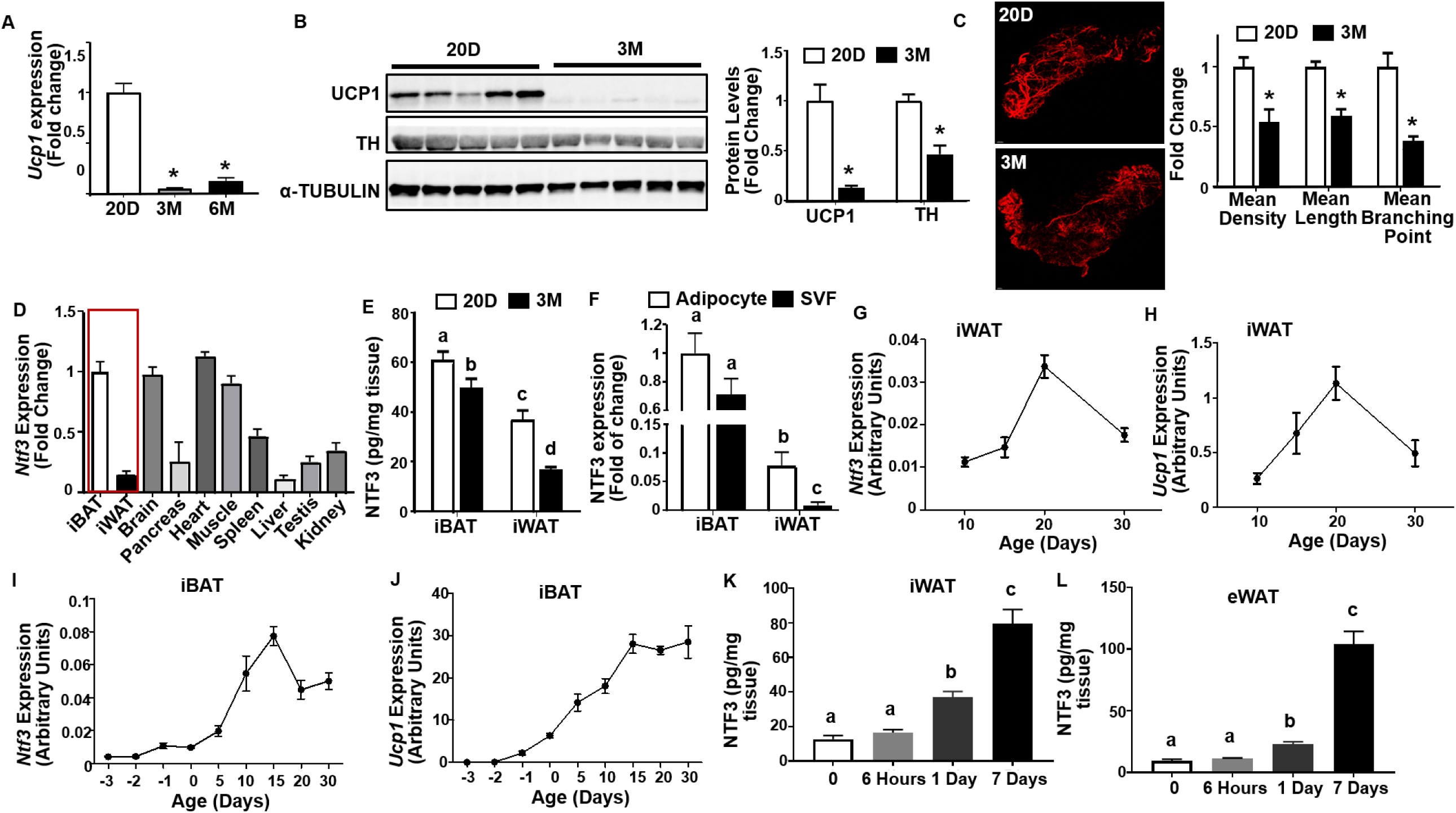
Adipose-derived neurotrophic factor NTF3 correlates with UCP1 expression and sympathetic innervation in adipose tissue. (A) *Ucp1* mRNA expression in iWAT of 20 days old postnatal pups, and 3- and 6-month-old mice (n=5). (B) UCP1 and TH protein levels in iWAT of 20 days old postnatal pups and 3 months old adult mice (n=5). (C) Representative images of TH-positive sympathetic nerve innervation in iWAT (left panel) and quantitation of mean nerve fiber density, mean nerve fiber length, and mean branching points normalized to total adipose tissue area (right panel) in iWAT of 20 days old postnatal pups and 3-month-old adult mice (n=3). (D) Tissue distribution of *Ntf3* expression (iBAT n=7; iWAT n=5; others n=4). (E) NTF3 protein levels measured by ELISA in iBAT and iWAT of 20 days old postnatal pups and 3 months old adult mice (BAT 20d n=6, 3M n=5; iWAT 20d n=10, 3M n=6). (F) The expression of *Ntf3* mRNA in adipocytes and stromal vascular fraction (SVF) in iBAT and iWAT (iBAT n=6; iWAT adipocyte n=8, SVF n=5). (G)-(H) The expression of *Ntf3* (G) and *Ucp1* (H) in iWAT during postnatal development in mice (n=4). (I)-(J) The expression of *Ntf3* (I) and *Ucp1* (J) in iBAT during postnatal development in mice (n=4). (K)-(L) NTF3 protein levels measured by ELISA in iWAT (K) and eWAT (L) during cold exposure (Time 0 n=4; 6h, 1d and 7d n=5). All data are expressed as mean ± SEM. Values that do not share a common superscript are significantly different at P < 0.05.

On the other hand, UCP1 and TH protein in iBAT was only slightly decreased in 3-month-old adult mice compared to that of P20 pups (**Suppl. Fig 1C**), suggesting that unlike WAT, UCP1 expression and sympathetic innervation in adult iBAT is largely preserved.

Existing literature supports an important role of neurotropic factor NTF3 in the regulation of normal SNS neuron function and growth and target tissue innervation ^23, 24^. We surveyed the expression patterns of neurotropic factors across mouse tissues. We found *Ntf3* was ubiquitously expressed in many major tissues, among which iBAT was one of the tissues with highest *Ntf3* expression; whereas *Ntf3* expression in iWAT was much lower (**Fig 1D**). In contrast, the expression of *Ngf, Bdnf* and *Nt4/5* was generally low and largely comparable in iBAT and iWAT (**Suppl. Fig 2**). Consistent with the RNA expression, NTF3 protein level was also much higher in iBAT than in iWAT, and exhibited significant decrease in both iBAT and iWAT of 3-month-old mice compared to that of P20 pups; the decrease was more profound in iWAT (**Fig 1E**). In addition, when measured in isolated adipocytes and stromal vascular fraction (SVF) in iBAT and iWAT, *Ntf3* was primarily expressed in mature adipocytes in iWAT; whereas *Ntf3* expression was comparable between adipocytes and SVF cells in iBAT (**Fig 1F**). Nonetheless, *Ntf3* expression was much higher in both adipocytes or SVF in iBAT than those in iWAT (**Fig 1F**).

Interestingly, the expression pattern of *Ntf3* in iWAT during postnatal development (**Fig 1G**) mimicked that of *Ucp1* (**Fig 1H**), peaks at P20, and then gradually decreased. The expression of *Ntf3* in iBAT peaked at 15 days of age and remained at a relatively high level (**Fig 1I**), similar to that of *Ucp1* expression pattern in iBAT (**Fig 1J**). In addition, cold exposure in adult mice significantly induced NTF3 protein levels in both iWAT and eWAT in a time-dependent manner (**Fig 1K** and **1L**). Our data suggest that NTF3 is important in maintaining BAT and WAT SNS innervation and *Ucp1* expression, and diminished NTF3 signaling in WAT may be responsible for the disappearance of beige adipocytes in adult mice via down-regulating SNS activity and innervation in WAT.

### Administration of NTF3 protein promotes beiging in postnatal mice and protects adult mice from diet-induced obesity

We reasoned that if fat tissue NTF3 level was maintained during early postnatal development, adequate SNS innervation in WAT might be maintained, which could also maintain adequate beige adipocyte phenotype. Indeed, intraperitoneal (i.p.) injection of NTF3 (50µg/kg body weight) daily in P20 mice for 10 days from P20 to P30 significantly increased *Ucp1* and other BAT-specific gene expression (**Fig 2A**), UCP1 and TH protein levels (**Fig 2B**) and UCP1-positive beige adipocytes in iWAT (**Fig 2C**); whereas NTF3 administration had minimal effects on *Ucp1* expression or UCP1 immunostaining in iBAT (**Suppl. Fig 3A-B**), which might be expected due to high endogenous NTF3 level in iBAT. These data suggest that continuous activation of NTF3 signaling in iWAT may be important in maintaining beige cell phenotype, and preventing them from disappearing during postnatal development.

**Figure 2.**
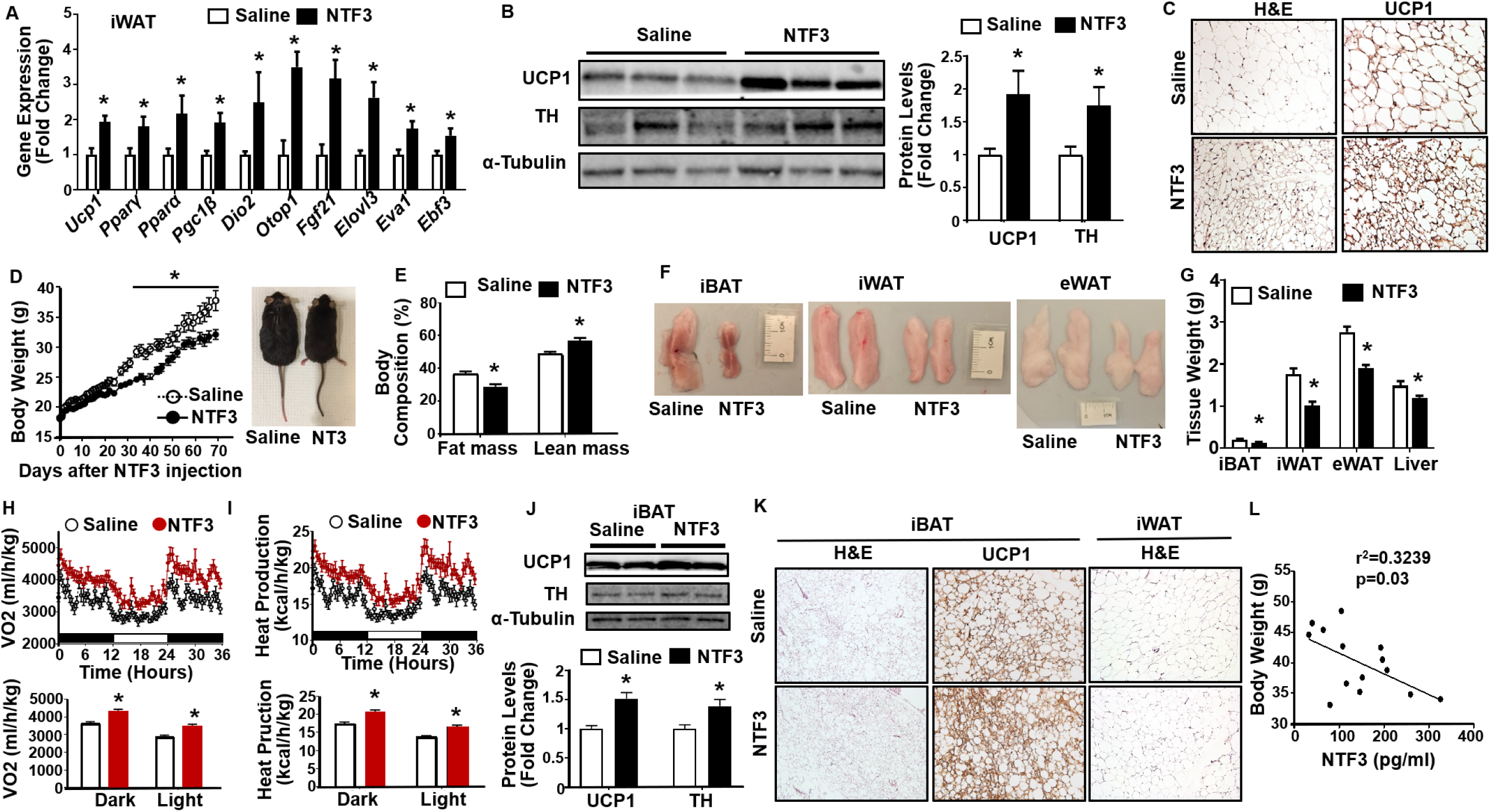
Administration of NTF3 protein promotes beiging in postnatal mice and protects adult mice from diet-induced obesity. (A)-(C) *Ucp1* and other BAT-specific gene expression (A) (n=5), UCP1 and TH protein levels (B) (n=4) and representative UCP1-immunostaining images showing UCP1-positive beige adipocytes in iWAT (C)(n=3) of 20 days old postnatal pups with daily intraperitoneal (IP) NTF3 (50 μg/kg) injection for 10 days (n=8). (D)-(G) Body weight (D), Body composition (E), Fat pad morphology (F) and Fat pad mass (G) in 6 weeks old mice with NTF3 (50µg/kg) ip injection every day for the first two weeks, then 3 times a week for 8 weeks while fed a HFD (saline n=9; NTF3 n=8). (H)-(I) Oxygen consumption (H) and Heat production (I) in 6 weeks old mice with NTF3 (50µg/kg) ip injection every day for the first two weeks, then 3 times a week for 8 weeks while fed a HFD (n=7). (J)-(K) UCP1 and TH protein levels in iBAT (J) (saline n=9, NTF3 n=8) and representative H&E and UCP1 immunostaining in iBAT and iWAT (K)(n=3) of 6 weeks old mice with NTF3 (50µg/kg) ip injection every day for the first two weeks, then 3 times a week for 8 weeks while fed a HFD. (L) Correlation between serum NTF3 levels and body weight in mice fed a HFD (n=14). All data are expressed as mean ± SEM. *p<0.05.

To further study the physiological significance of NTF3 in regulating energy homeostasis in adult animals, we injected NTF3 (50µg/kg body weight) into 6-week-old C57BL/6J mice fed with a high fat diet (HFD). As expected, serum NTF3 level was significantly increased in mice with NTF3 injection (**Suppl. Fig 4A**). Mice injected with NTF3 were protected from diet-induced obesity as they gained less weight on HFD starting around 5 weeks after NTF3 treatment (**Fig 2D**). The NTF3-injected mice showed reduced fat mass with concomitantly increased lean mass (**Fig 2E**). In consistence, we observed decreased fat pad mass in iWAT, eWAT and iBAT depots, and decreased liver weight in NTF3-injected mice (**Fig 2F-G**). Mice with NTF3 injection had higher oxygen consumption and heat production compared to saline-injected mice (**Fig 2H-I**), whereas there was no change in respiratory exchange rate (RER), locomotor activity and food intake (**Suppl. Fig 4B-D**). These data suggest that the reduced adiposity in NTF3-injected mice may be primarily caused by increased energy expenditure. In consistence, we discovered enhanced UCP1 and TH protein expression in iBAT of NTF3-injected mice measured by immunoblotting (**Fig 2J**), more UCP1 immunostaining in iBAT (**Fig 2K**), and smaller adipocytes in both iBAT and iWAT (**Fig 2K**). Mice with NTF3 injection also exhibited lower fed glucose level (**Suppl. Fig 4E**) and improved insulin sensitivity in insulin tolerance (ITT) test (**Suppl. Fig 4F**). In addition, there was an inverse correlation between serum NTF3 levels and body weight in mice fed with HFD (**Fig 2L**). Thus, our data strongly suggest that NTF3 injection protects mice from diet-induced obesity, primarily due to increased energy expenditure.

### Overexpression of Ntf3 in adipocytes promotes cold-induced thermogenesis and protects mice from diet-induced obesity

We have generated a mouse model with NTF3 overexpression specifically in adipocytes under the control of the adiponectin promoter (AdNTF3-Tg) (**Suppl. 5A-B**), which resulted in increased *Ntf3* mRNA levels in iWAT (**Fig 3A**), and increased NTF3 protein levels in iWAT as measured by both immunoblotting (**Fig 3B**) and ELISA (**Fig 3C**). AdNTF3-Tg mice also exhibited increased NTF3 protein levels in eWAT and to a lesser extent in iBAT, without changes of NTF3 protein levels in circulation (**Suppl. Fig 5C-E**). We therefore established an ideal transgenic mouse for the study of the direct effect of the fat-derived NTF3 in sympathetic innervation, which is unlikely confounded by the systemic effect of NTF3 due to the restriction of *Ntf3* overexpression to the local adipose tissue.

**Figure 3.**
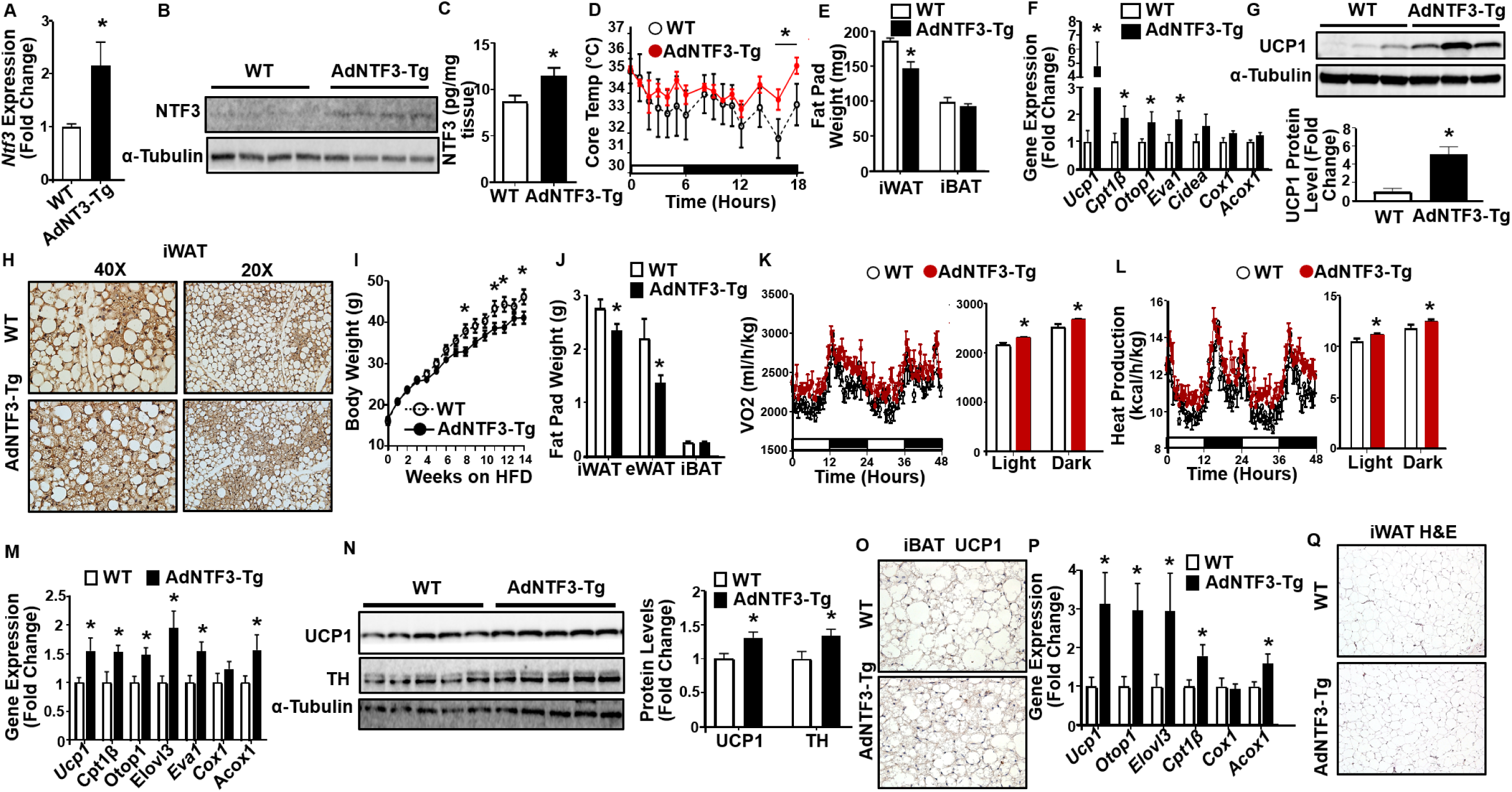
Overexpression of NTF3 in adipocytes promotes cold-induced thermogenesis and protects mice from diet-induced obesity. (A)-(C) *Ntf3* mRNA (A) (n=3), NTF3 protein levels measured by immunoblotting (B) (WT n=7, AdNTF3-Tg n=5) and ELISA (C) (WT n=3, AdNTF3-Tg n=4) in iWAT of WT and AdNTF3-Tg mice. (D)-(H) Core body temperature (D)(n=4), Fat pad mass (E)(n=4), *Ucp1* and other thermogenic gene expression in iWAT (F)(n=4), UCP1 protein levels in iWAT (G)(n=3), and representative UCP1 immunostaining images showing UCP1 positive beige adipocytes in iWAT (H)(n=3) of AdNTF3-Tg mice and their littermates during cold exposure. (I)-(L) Body weight (I) (WT n=8, AdNTF3-Tg n=10), Fat pad mass (J) (WT n=6, AdNTF3-Tg n=7), Oxygen consumption (K) (WT n=5, AdNTF3-Tg n=6) and Heat production (L) (WT n=5, AdNTF3-Tg n=6) in AdNTF3-Tg mice and their littermate controls fed with a HFD starting 6 weeks of age. (M)-(O) *Ucp1* and other thermogenic gene expression (M) (n=7), UCP1 and TH protein levels (N) (n=8), and representative UCP1 immunostaining in iBAT (O)(n=3) of AdNTF3-Tg mice and their littermate controls fed a HFD. (P)-(Q) *Ucp1* and other thermogenic gene expression (P) (n=6), and H&E staining (Q)(n=3) in iWAT of AdNTF3-Tg mice and their littermate controls fed a HFD. All data are expressed as mean ± SEM. *p<0.05.

We first assessed cold-induced thermogenesis in AdNTF3-Tg mice. Two months old AdNTF3-Tg mice and their wildtype (WT) littermate controls were subjected to a cold challenge (5°C) for 7 days. Interestingly, AdNTF3-Tg mice exhibited a tendency of higher body temperature during cold exposure compared to WT, which reached significance at 16-18 hours after the start of cold exposure (**Fig 3D**), suggesting cold resistance in these transgenic mice. After a 7-day cold exposure, AdNTF3-Tg mice displayed lower fat pad mass in iWAT (**Fig 3E**), enhanced expression of thermogenic genes such as *Ucp1, Cpt1β, Otop1* and *Eva1* in iWAT (**Fig 3F**), increased UCP1 protein expression (**Fig 3G**) and appearance of more UCP1-positive multilocular cells in iWAT (**Fig 3H**), indicating an increased beigeing phenotype in AdNTF3-Tg mice in response to a chronic cold challenge.

Overexpressing NTF3 in adipocytes also led to increased thermogenic gene expression in iBAT, including *Ucp1, Pparγ, Cpt1β* and *Eva1* (**Suppl. Fig 5F**). UCP1 protein tended to increase in iBAT of AdNTF3-Tg mice after the 7-day cold exposure (p=0.07, **Suppl. Fig 5G**). Histological analysis from both H&E staining and UCP1-immunostaining showed decreased lipid droplet sizes in iBAT of AdNTF3-Tg mice compared to that of WT mice after cold exposure (**Suppl. Fig 5H**). Our data indicate that NTF3 overexpression in adipocytes results in overall improvement of brown and beige adipocyte thermogenic function against a cold challenge.

We then challenged AdNTF3-Tg and WT mice with HFD by weaning them onto the HFD. Body weight of the AdNTF3-Tg and WT mice started diverging after 6-8 weeks of HFD feeding, with the AdNTF3-Tg mice gaining less weight (**Fig 3I**). This was associated with decreased fat pad mass in iWAT and eWAT of the transgenic mice (**Fig 3J**). The reduced adiposity in the AdNTF3-Tg mice might stem from increased energy expenditure, as they exhibited increased oxygen consumption and heat production (**Fig 3K-L**) without changes in food intake and locomotor activity (**Suppl. Fig 6A-B**). In consistence, AdNTF3-Tg mice exhibited enhanced expression of BAT-specific genes such as *Ucp1, Cpt1b, Otop1, Elovl3, Eva1*, and *Acox1* in iBAT (**Fig 3M**), increased UCP1 and TH protein levels in iBAT as measured by immunoblotting (**Fig 3N**), and smaller brown adipocytes with more UCP1 immunostaining in iBAT compared to that of WT controls (**Fig 3O**). Moreover, AdNTF3-Tg mice had increased expression of thermogenic genes including *Ucp1, Otop1, Eovl3, Cpt1b* and *Acox1* in iWAT (**Fig 3P**), and reduced adipocyte size in iWAT as shown by H&E staining compared to that of WT mice (**Fig 3Q**). With decreased body weight and adiposity, these transgenic mice exhibited increased insulin sensitivity in glucose tolerance (GTT) and ITT tests (**Suppl. Fig 6C-D**). Our data suggest that the enhanced thermogenic program in iBAT and iWAT collectively contributes to the increased energy expenditure in transgenic mice with adipocyte overexpression of NTF3, resulting in reduced adiposity and improved insulin sensitivity when challenged with HFD feeding.

To test the reproducibility of the genetic model with *Ntf3* overexpression, we have also generated mice that overexpress *Ntf3* specifically in adipocytes by inserting the *Ntf3* transgene with loxP-flanked Stop sequence into the *Rosa26* locus (**Suppl. Fig 7A-B**) as previously described ^25^. The resulting Rosa-NTF3^fl/+^ mice were crossed with Adiponectin-Cre mice ^26^ (Kindly provided by Dr. Evan Rosen, Beth Israel Medical Center, Harvard Medical School) to generate mice with adipocyte-specific NTF3 knock-in mice (Adiponectin-Cre::Rosa-NTF3^fl/+^, or AdNT3-KI) by deleting the loxP-flanked Stop sequence. AdNTF3-KI mice had significantly increased *Ntf3* mRNA and protein levels in BAT and WAT compared to fl/+ controls (**Supp. Fig 7C-D**). However, unlike our AdNTF3-Tg mice, where NTF3 overexpression was primarily restricted within adipose tissue (**Fig 3A-C, Suppl. Fig 5C-E**), AdNTF3-KI mice exhibited significantly increased circulating NTF3 levels compared to fl/+ controls (**Suppl. Fig 7E**).

Similar to AdNTF3-Tg mice, AdNTF3-KI mice exhibited enhanced cold-induced thermogenesis as shown by higher body temperature, lower body weight, lower fat mass, increased expression of *Ucp1* and other thermogenic genes in iWAT, higher UCP1 and TH protein levels in iWAT and more UCP1-positive beige adipocytes in iWAT than those of fl/+ control mice in response to a 7-day cold challenge (**Suppl. Fig 7F-K**).

In consistence, AdNTF3-KI mice had lower body weight, lower body fat content, and reduced fat pad mass in iBAT, iWAT and rWAT than fl/+ controls when challenged with a HFD (**Suppl. Fig 8A-C**). The reduced adiposity in AdNTF3-KI mice was primarily due to increased energy expenditure, as AdNF3-KI mice had no difference in food intake (**Suppl. Fig 8D**), but had significantly increased oxygen consumption and heat production (**Suppl. Fig 8E-F**), without changes in locomotor activity (**Suppl. Fig 8G)**. Consistent with their higher energy expenditure, AdNTF3-KI mice exhibited increased *Ucp1* and other thermogenic gene expression, increased UCP1 and TH protein levels, and smaller brown adipocyte size with more intense UCP1 immunostaining in iBAT (**Suppl. Fig 8H-J**). Because of their lean phenotype, AdNT3-KI mice were more insulin sensitive compared to fl/+ controls during the HFD challenge as assessed by GTT and ITT tests (**Suppl. Fig8K-L**).

### NTF3 regulates sympathetic ganglia (SG) neuronal growth and activation

We then further studied mechanisms by which adipose tissue-derived NTF3 regulates brown and beige adipocyte thermogenic function. Since NTF3 is a neurotropic factor that regulates SNS neuron growth and target tissue innervation ^23, 24^, we first tested NTF3’s effect on sympathetic neurite growth in primarily cultured sympathetic neurons *in vitro*. We found that NTF3 (100ng/ml) treatment significantly stimulated sympathetic neuron neurite growth, as shown by increased number of neurites per neuron, total neurite length and maximal neurite length per neuron, as well as average neurite length per neurite per neuron (**Fig 4A**).

**Figure 4.**
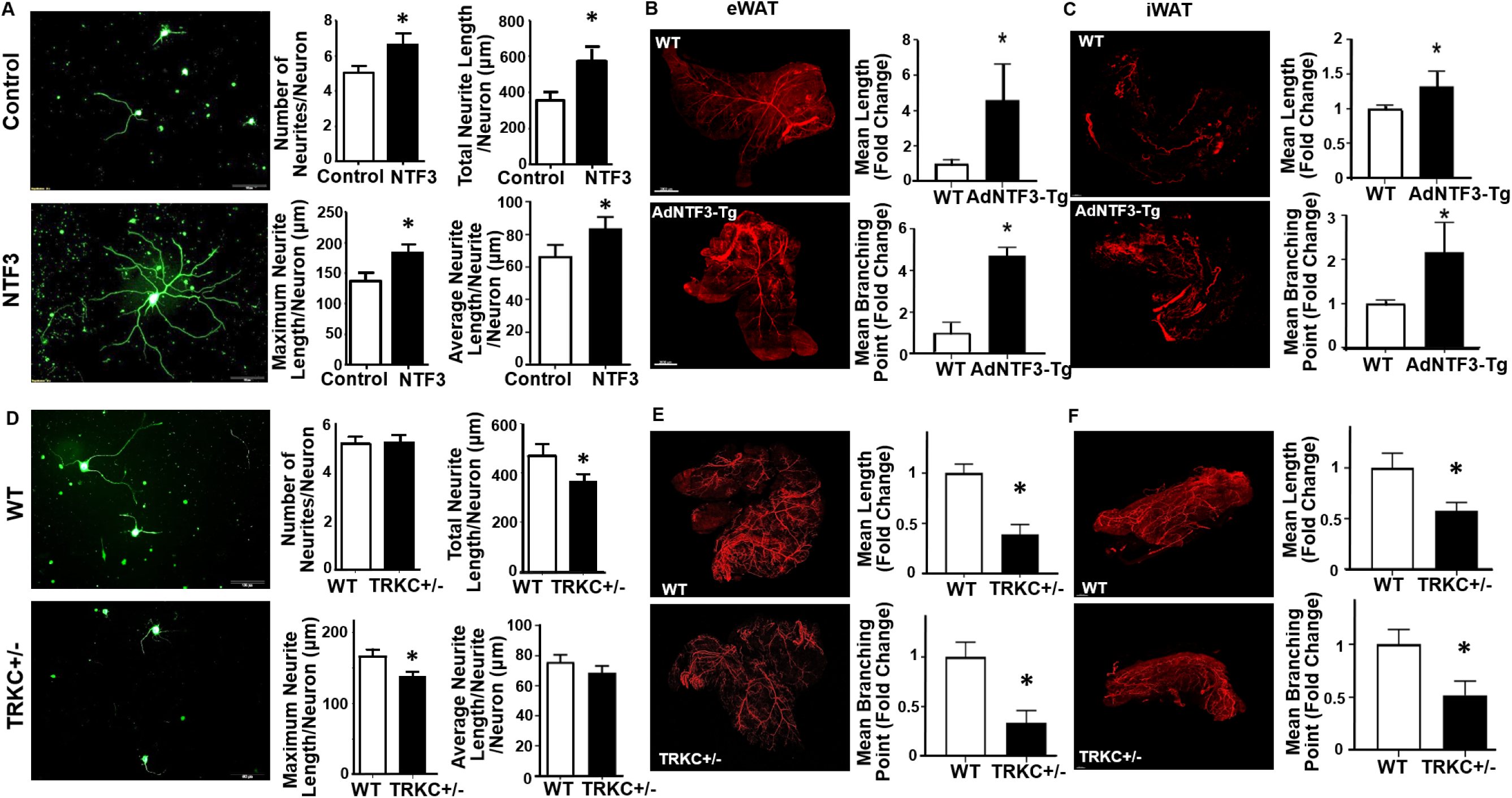
NTF3/TRKC regulates sympathetic neuronal growth and innervation in adipose tissue. (A) Representative βIII-tubulin immunofluorescent images of sympathetic ganglia neurons treated with NTF3 (100ng/ml) (left panel) and quantitation of number of neurites per neuron, total neurite length per neuron, maximal neurite length per neuron and average neurite length per neurite per neuron (right panel) (Control n=80, NTF3 n=60). (B)-(C) Representative images of eWAT (B) and iWAT (C) TH-positive sympathetic nerve innervation (left panel) and quantitation of mean nerve fiber length and mean nerve fiber branching points normalized to total adipose tissue area (right panel) in AdNT3-Tg mice and their littermate control mice after a 7-day cold challenge (n=3). (D) Representative βIII-tubulin immunofluorescent images of sympathetic ganglia neurons isolated from WT and TRKC+/- mice (left panel) and quantitation of number of neurites per neuron, total neurite length per neuron, maximal neurite length per neuron and average neurite length per neurite per neuron (right panel) (control n=158, TRKC+/- n=162). (E)-(F) Representative images of eWAT (E) and iWAT (F) TH-positive sympathetic nerve innervation (left panel) and quantitation of mean nerve fiber length and mean nerve fiber branching points normalized to total adipose tissue area (right panel) in TRKC+/- mice and their littermate control mice after a 7-day cold challenge (n=3). All data are expressed as mean ± SEM. *p<0.05.

The overexpression of NTF3 in our AdNTF3-Tg model is primarily restricted to the adipose tissue, without any confounding effects on the changes in the circulating NTF3. Thus, to further assess the effect of adipose tissue NTF3 on sympathetic nerve innervation into adipose tissue *in vivo*, we immunostained sympathetic nerves innervating adipose tissue with antibodies against sympathetic marker TH using the Adipo-Clear approach ^22^ in AdNTF3-Tg mice. Analysis of the sympathetic nerve fibers by Imaris Image Analysis Software revealed a significantly higher mean nerve fiber length and mean nerve fiber branching points in iWAT of 2-month-old AdNTF3-Tg mice than that in WT mice when housed at ambient room temperature (**Suppl. Fig 9**). In addition, a 7-day cold exposure further increased mean nerve fiber length and nerve fiber branching points in eWAT and iWAT of 2-month-old AdNTF3-Tg mice compared to that of WT mice (**Fig 4B-C**). Our data strongly support an important role of adipose tissue-derived NTF3 in regulating sympathetic nerve innervation in fat tissues, which may contribute to its effects on promoting brown and beige adipocyte thermogenic function during cold and diet challenges as observed in NTF3-injected, AdNTF3-Tg and AdNTF3-KI mice (**Fig 2-3, Suppl. Fig 7-8**).

### NTF3/TRKC regulates sympathetic innervation in adipose tissue

NTF3 binds with high affinity to its cognitive receptor, neurotrophic receptor 3/Tropomyosin receptor kinase C (NTRK3/TRKC) ^27, 28, 29^. TRKC immunoreactivity has been found in majority of sympathetic neurons in SG during embryonic development and in adult animals ^30, 31^, indicating the importance of TRKC signaling in regulating sympathetic neuron growth and development.

To further explore mechanisms by which NTF3 regulates sympathetic innervation in adipose tissue, we investigated whether TRKC, the receptor for NTF3, is important in regulating sympathetic innervation in adipose tissue. Homozygous TRKC-/- mice die by postnatal day 21 ^32^. Thus, we studied mice with haploinsufficiency of TRKC (TRKC+/-). As expected, TRKC expression in SG was reduced to approximately 50% in TRKC+/- mice compared to that of WT littermates (**Suppl. Fig 10**). Interestingly, sympathetic neurons isolated from TRKC+/- mice had significantly suppressed neuronal growth, as shown by decreased total neurite length and maximal neurite length per neuron (**Fig 4D**).

Using Adipo-Clear approach with TH immunostaining, we found that TRKC+/- mice had significantly less sympathetic innervation in adipose tissue in response to a 7-day cold challenge when compared to that of WT control mice, as shown by reduced mean nerve fiber length and mean nerve fiber branching points in eWAT and iWAT of TRKC+/- (**Fig 4E-F**). Our data strongly support the importance of NTF3/TRKC in regulating sympathetic innervation in adipose tissue.

### NTF3/TRKC regulates SG neuronal activity

To further study whether NTF3/TRKC regulates the activity of SG that specifically innervate iBAT, we injected fluorescent retrograde neuronal tracer Fast Blue (FB) into iBAT of TRKC-Cre::R26-stop-EYFP reporter mice. Animals were given 10-14 days to recover and allow for retrograde FB transport, and were then subjected to a 7-day cold challenge. SG at the thoracic T2 level, which innervates iBAT ^33^, were analyzed for single-, double- and triple-labeled neurons (**Fig 5A-C**). Interestingly, the majority of the TRKC-positive neurons in T2 SG were co-localized with TH (TH^+^TRKC^+^) (**Fig 5A-B**). In addition, approximately 17% of TH^+^TRKC^+^ neurons in mice housed at room temperature were also co-localized with FB, indicating that these neurons specifically innervate iBAT (**Fig 5C**), with the remaining 83% of FB^-^ neurons in TH^+^TRKC^+^ populations in the T2 SG innervate organs other than iBAT (**Fig 5C**). Cold exposure significantly increased the percentage of TH^+^TRKC^+^FB^+^ neurons in T2 ganglia (**Fig 5B-C**), with concomitant decreases in the percentage of TH^+^ single-labeled and TH^+^FB^+^ double labeled neurons; whereas there was no significant change in the percentage of TH^+^TRKC^+^FB^-^ neurons (**Fig 5B-C**). Our data indicate that cold exposure specifically increases TH^+^TRKC^+^ neurons in the SG that innervate brown fat (TH^+^TRKC^+^FB^+^ neurons), whereas it does not affect TH^+^TRKC^+^ neurons that might innervate organs other than iBAT (TH^+^TRKC^+^FB^-^ neurons).

**Figure 5.**
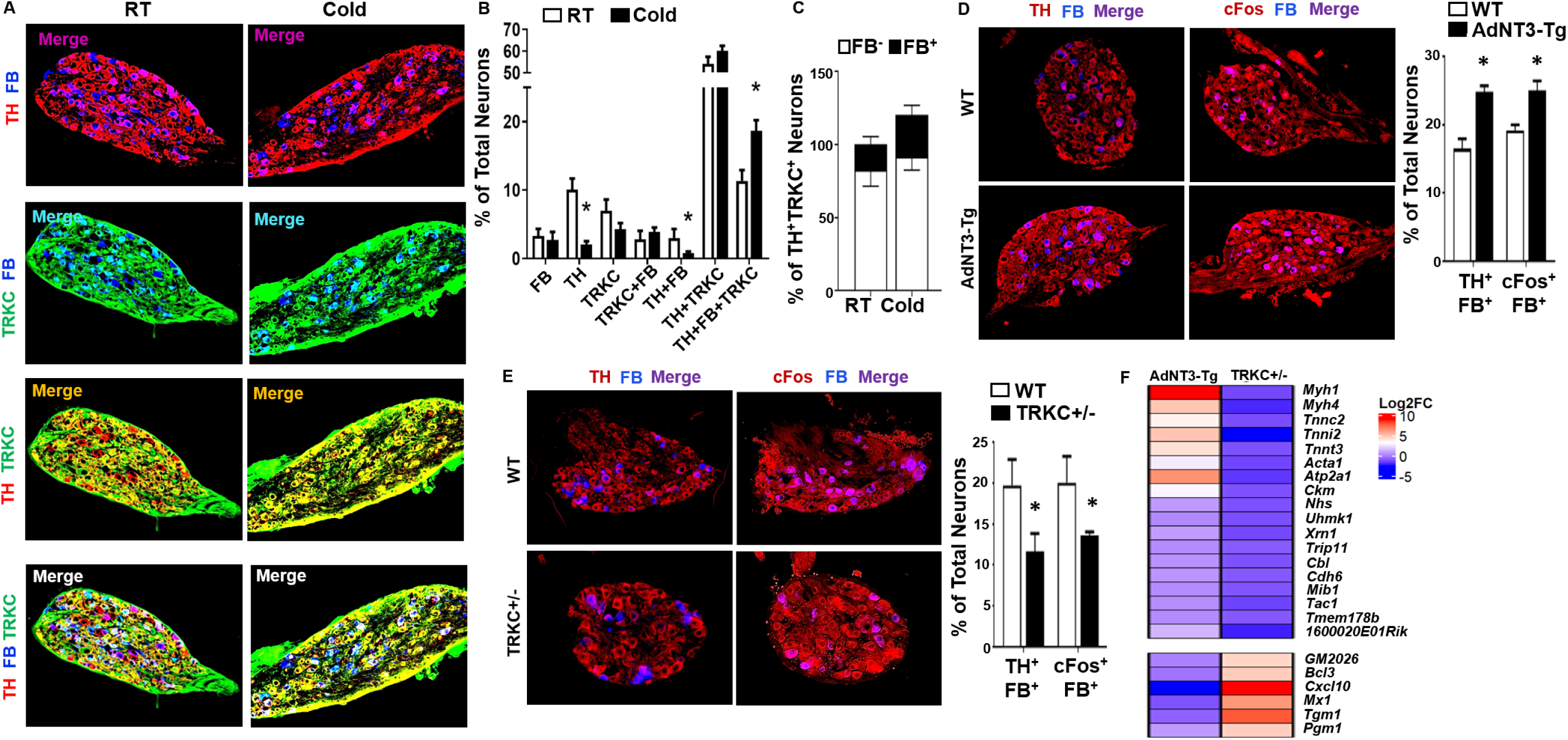
NTF3/TRKC regulates sympathetic ganglia neuronal activity. (A)-(B) Representative images of TH, Fast Blue (FB) and TRKC-EYFP labeling (A), and percentage of single-, double- and triple-labeled neurons (B) in sympathetic ganglia at thoracic T2 level in TRKC-Cre::R26EYFP mice at room temperature (RT) or subjected to a 7-day cold challenge (RT n=5, COLD n=7). (C) Percentage of FB-positive (FB+) and FB-negative (FB-) neurons in TH/TRKC-positive neurons in sympathetic ganglia at thoracic T2 level in TRKC-Cre::R26EYFP mice at room temperature (RT) or subjected to a 7-day cold challenge (RT n=5, COLD n=7). (D) Representative images (left panel) and quantitation (right panel) of TH/FB and cFos/FB labeling in sympathetic ganglia at thoracic T2 level in WT and AdNTF3-Tg mice after a 7-day cold challenge (n=4). (E) Representative images (left panel) and quantitation (right panel) of TH/FB and cFos/FB labeling in sympathetic ganglia at thoracic T2 level in WT and TRKC+/- mice after a 7-day cold challenge (n=4). (F) Heatmap of the expression of genes regulated in the opposite direction in sympathetic ganglia from thoracic T1 to T4 levels of AdNTF3-Tg and TRKC+/- mice after 1-day cold challenge. T1-T4 sympathetic ganglia from 6 mice each group were pooled for RNAseq analysis. All data are expressed as mean ± SEM. *p<0.05.

We then injected FB into iBAT of AdNTF3-Tg, TRKC+/- and their respective control mice and subjected them to a 7-day cold challenge. SG at thoracic T2 level were collected for immunohistochemistry analysis with antibodies against TH as the sympathetic marker or cFos as the neuronal activation marker, and double-stained with FB. Interestingly, overexpressing NTF3 in adipose tissue significantly increased, while TRKC haploinsufficiency significantly decreased TH/FB or cFos/FB double-labeled neurons in T2 SG (**Fig 5D-E**) during cold exposure.

We have similarly injected FB into iWAT of AdNTF3-Tg, TRKC+/- and their respective littermate control mice and subjected them to a 7-day cold challenge. SG at lumbar L1 level, which innervate iWAT ^33^, were collected for immunohistochemistry analysis for TH/FB or cFos/FB immunostaining. We found that TH/FB or cFos/FB double-labeled neurons were significantly increased in L1 SG of AdNTF3-Tg mice; but significantly decreased in L1 SG of TRKC+/- mice when compared to their respective littermate controls (**Suppl. Fig 11A-B**).

Consistent with reduced sympathetic innervation and activity in TRKC+/- mice, basal norepinephrine (NE) level was reduced in fat tissues including iWAT, eWAT and rWAT after a 16-hour cold exposure in TRKC+/- mice (**Suppl. Fig 12A**). In addition, NE turnover rate (NETO) was also significantly reduced in all fat tissues including iBAT, iWAT, eWAT and rWAT after cold exposure in TRKC+/- mice (**Supple. Fig 12B**).

Thus, our data strongly support the importance of adipose tissue derived NTF3 and TRKC in the regulation of peripheral sympathetic neuron activity in SG that directly innervate brown and white adipose tissue.

### NT3/TRKC regulates a plethora of transcriptional program that controls neuronal axonal growth and elongation in SG

To gain further insight into how NT3/TRKC regulates sympathetic neuronal function and axonal growth, we performed RNAseq analysis in thoracic T1-T4 SG that innervate iBAT ^33^ from 1-day cold-challenged AdNTF3-Tg, TRKC+/- and their respective WT control mice.

Our RNAseq analysis demonstrated that pathways involved in neuronal axonal growth were regulated in an opposite direction in SG of AdNTF3-Tg and TRKC+/- mice (**Fig 5F**). For example, axonal outgrowth and guidance to the proper target tissues requires the coordination of cellular programs, including the coordination of actin filaments and microtubules, the dynamic cytoskeletal polymers and “building blocks” that promote shape change and locomotion during the peripheral nerve axonal outgrowth and elongation ^34, 35^. Interestingly, we found several myosins, actins and troponins were upregulated in AdNTF3-Tg mice while downregulated in TRKC+/- mice, including myosin heavy polypeptide 1 (*Myh1*), myosin heavy polypeptide 4 (*Myh4*), actin alpha 1 (*Acta1*), troponin C2 (*Tnnc2*), troponin I skeletal, fast 2 (*Tnni2*), troponin T2 (*Tnnt2*), muscle creatine kinase (*Ckm*), and sarcoplasmic/endoplasmic reticulum calcium ATPase 1 (*Atp2a1*) (**Fig 5F**). Although these genes are typically characteristic of striated muscle contraction pathway, similar genes in this pathway, including *Myh1* and *ATP2a1*, were shown to be decreased in the brain of the neuron degenerative disease Amyotrophic lateral sclerosis (ALS) ^36^. Thus, these myosin and actin filament components may be novel members of the cytoskeletal family regulated by NTF3/TRKC signaling important for axonal growth and elongation in SG.

In addition, recent evidence also supports the axonal transport and translation of specific mRNAs as a mechanism of quickly localizing needed proteins at the axons and axonal growth cones ^34, 35^. For example, β-actin mRNA was localized to axons and axonal growth cones, and this localization can be induced by NTF3 signaling; blocking this process also blocked NTF3-induced protein localization at the growth cone and decreased growth cone motility ^37, 38^. Here, we found that the serine/threonine kinase *Uhmk1* and the 5’-3’ exonuclease *Xrn1*, genes that are involved in synaptic mRNA transport and translation ^39, 40^, were upregulated in AdNTF3-Tg mice but downregulated in TRKC+/- mice (**Fig 5F**).

Further, the multifunctional adaptor protein casitas B-lineage lymphoma (*Cbl*) has been shown to be necessary for the recruitment of NGF/TRKA to the lipid raft and mediates signals important for actin reorganization and neurite growth ^41, 42^. Interestingly, we found that *Cbl* was upregulated in AdNTF3-Tg mice but downregulated in TRKC+/- mice (**Fig 5F**), indicating *Cbl* may also be regulated by NTF3/TRKC signaling.

Additional genes that were oppositely regulated in AdNTF3-Tg and TRKC+/- mice include proteins that regulate cytoskeletal dynamics and activities (MX dynamin-like GTPase 1 (*Mx1*), transglutaminase 1, k polypeptide (*Tgm1*), and NHS actin remodeling regulator (*Nhs*)); proteins involved in glucose metabolism (phosphoglucomutase 1 (*Pgm1*); intracellular protein sorting (thyroid hormone receptor interactor 11 (*Trip11*), and mindbomb E3 ubiquitin protein ligase 1 (*Mib1*)) and the chemokine (c-x-c motif) ligand 10 (*Cxcl10*) (**Fig 5F**). Interestingly, it was reported that *Cxcl10* in the spinal cord was upregulated following spinal cord injury, which was critical in recruiting T lymphocytes and inducing apoptosis at the injury site; whereas neutralizing CXCL10 resulted in reduced apoptosis and increased axonal sprouting following spinal cord injury ^43, 44^. Thus, our data may identify a novel link between inflammation and neuronal axonal growth, which may be regulated by NTF3/TRKC signaling.

### Mice with haploinsufficiency of TRKC exhibit impaired cold-induced thermogenesis and are prone to diet-induced obesity

We then further studied the importance of TRKC in regulating cold- and diet-induced thermogenesis and whole body energy homeostasis. During a 7-day cold challenge, TRKC+/- mice exhibited lower body temperature (**Fig 6A**) and were therefore more cold sensitive than their littermate controls. During the cold challenge, TRKC+/- mice had significantly reduced *Ucp1* and other thermogenic gene expression in iWAT, retroperitoneal WAT (rWAT) and iBAT (**Fig 6B-D**), reduced UCP1 and TH protein levels in iWAT and rWAT (**Fig 6E-F**), and larger adipocytes but much fewer UCP1-positive multilocular beige adipocytes in iWAT and rWAT (**Fig 6G**). However, there was no significant difference in UCP1 protein levels and UCP1 immunostaining in iBAT between TRKC+/- and WT control mice during the 7-day chronic cold challenge (**Suppl. Fig 13A-B**). These data suggest that TRKC is important in regulating cold-induced thermogenesis, especially cold-induced beigeing in WAT.

**Figure 6.**
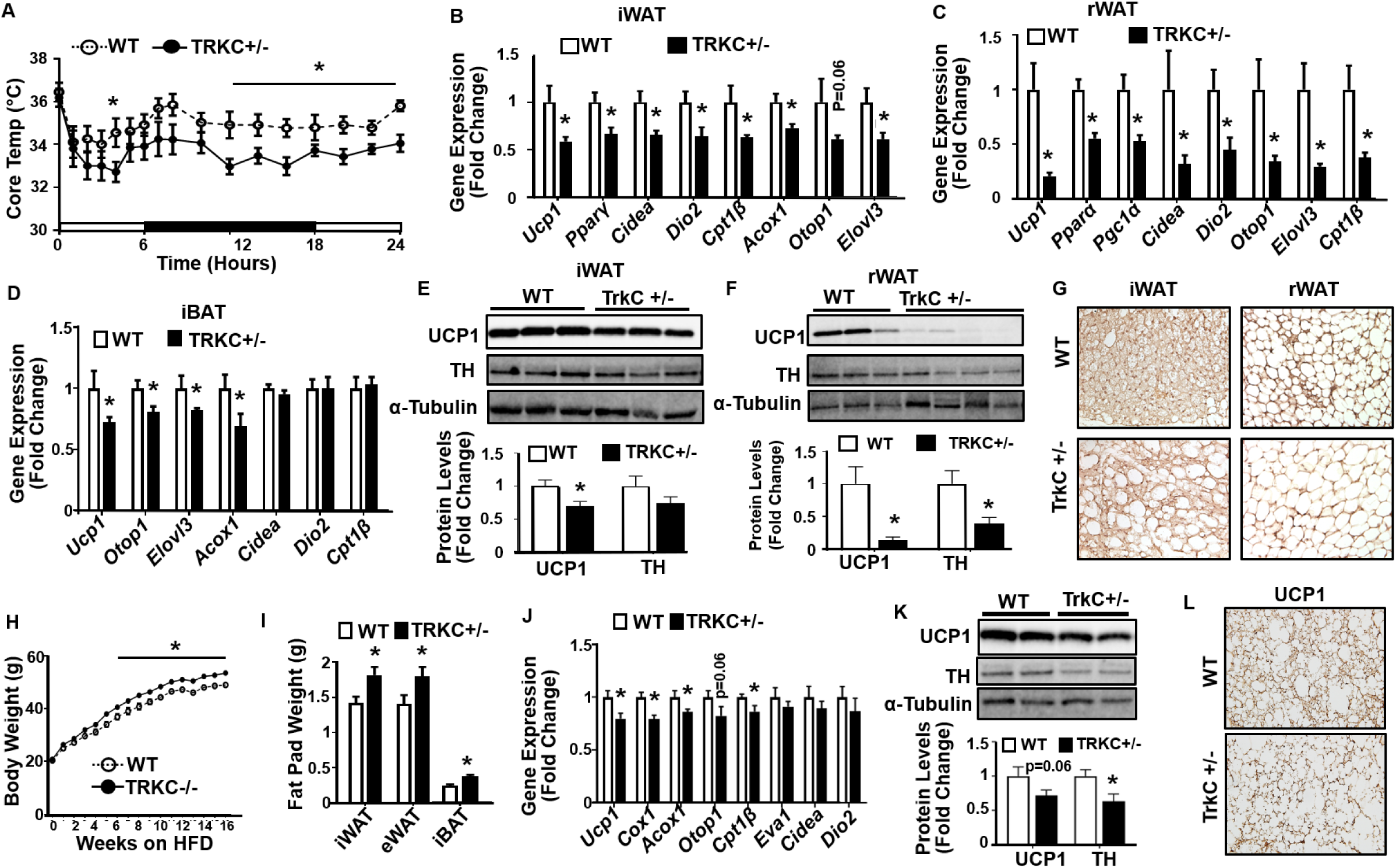
Mice with haploinsufficiency of TRKC exhibit impaired cold-induced thermogenesis and are prone to diet-induced obesity. (A)-(D) Body temperature (A) (WT n=8, TRKC+/- n=6), and *Ucp1* and other thermogenic gene expression in iWAT (B) (WT n=4, TRKC+/- n=5), rWAT (C) (n=5) and iBAT(D) (WT n=4, TRKC+/- n=5) of WT and TRKC+/- mice during a 7-day cold exposure (WT n=4, TRKC+/- n=5). (E)-(G) UCP1 and TH protein levels in iWAT (E)(WT n=5, TRKC+/- n=3) and rWAT (F)(WT n=3, TRKC+/- n=4), and representative UCP1 immunostaining images showing UCP1 positive beige adipocytes in iWAT and rWAT (G)(n=3) of WT and TRKC+/- mice during a 7-day cold exposure. (H)-(L) Body weight (H) (WT n=7, TRKC+/- n=8), Fat pad mass (I) (BAT, WT n=7, TRKC+/- n=8; iWAT, eWAT, rWAT, WT n=10, TRKC+/- n=14), *Ucp1* and other thermogenic gene expression in iBAT (J) (BAT WT n=8, TRKC+/- n=10), UCP1 and TH protein levels in iBAT (K) (WT n=6, TRKC+/- n=5), and representative UCP1 immunostaining in iBAT (L)(n=3) of WT and TRKC+/- mice fed a HFD under thermoneutrality (30°C). All data are expressed as mean ± SEM. *p<0.05.

We then subjected TRKC+/- mice with a HFD challenge. When housed at ambient room temperature (20-22°C), TRKC+/- mice on HFD had increased fat mass in eWAT and rWAT and had a tendency of increase in iWAT mass, despite no change in body weight and food intake (**Suppl. Fig 14A-C**). Interestingly, TRKC+/- mice exhibited decreased energy expenditure evident by reduced oxygen consumption and heat production (**Suppl. Fig 14D**-**E**) without changes in locomotor activity (**Suppl. Fig 14F**). The decreased energy expenditure in TRKC+/- mice was associated with a tendency of reduced UCP1 and significantly reduced TH protein levels in iBAT (**Suppl. Fig 14G**). Mice with haploinsufficiency of TRKC also showed glucose intolerance and insulin resistance in GTT and ITT tests, respectively (**Suppl. Fig 14H-I**).

We also performed HFD feeding in these animals under thermoneutrality (30°C) to avoid any non-shivering thermogenesis that may be induced by mild cold stress in mice housed under ambient room temperature (20-22°C) as we recently reported ^45^, which may serve as a confounding factor in assessing diet-induced thermogenesis ^4, 46, 47^. When housed at thermoneutrality, HFD-fed TRKC+/- mice consistently gained more weight (**Fig 6H**) and had increased fat mass in iBAT, iWAT and eWAT (**Fig 6I**) with larger adipocytes sizes (**Suppl. Fig 15A**) compared to their WT control mice. Moreover, TRKC+/- mice had down-regulated expression of thermogenic genes in iBAT (**Fig 6J**) and had a tendency of reduced UCP1 protein levels and significantly reduced TH protein levels in iBAT (**Fig 6K**). This was associated with less UCP1-immunostaining and enlarged brown adipocytes in iBAT (**Fig 6L**). TRKC+/- mice also exhibited reduced thermogenic gene expression in both iWAT and eWAT (**Suppl. Fig 15B-C**). Consistent with increased adiposity, TRKC+/- mice exhibited glucose intolerance and insulin resistance in GTT and ITT tests, respectively (**Suppl. Fig 15D-E**). In sum, our data suggest that TRKC signaling is crucial in regulating cold- and diet-induced thermogenesis and energy balance.

### Mice with specific deletion of TRKC in sympathetic nerves exhibit impaired cold-induced thermogenesis and are prone to diet-induced obesity

We have generated mice with sympathetic-specific deletion of TRKC by crossing TRKC-floxed mouse model (fl/fl) where the TRKC kinase ATP-binding domain is flanked with loxP sites ^48^ with TH-Cre mice where Cre-recombinase expression is under the control of the TH promoter ^49^ to achieve specific deletion of TRKC in sympathetic neurons (TH-Cre::TRKC-fl/fl, or STRKCKO). We found that TRKC expression was reduced around 75% in SG (**Fig 7A**) of STRKCKO mice.

**Figure 7.**
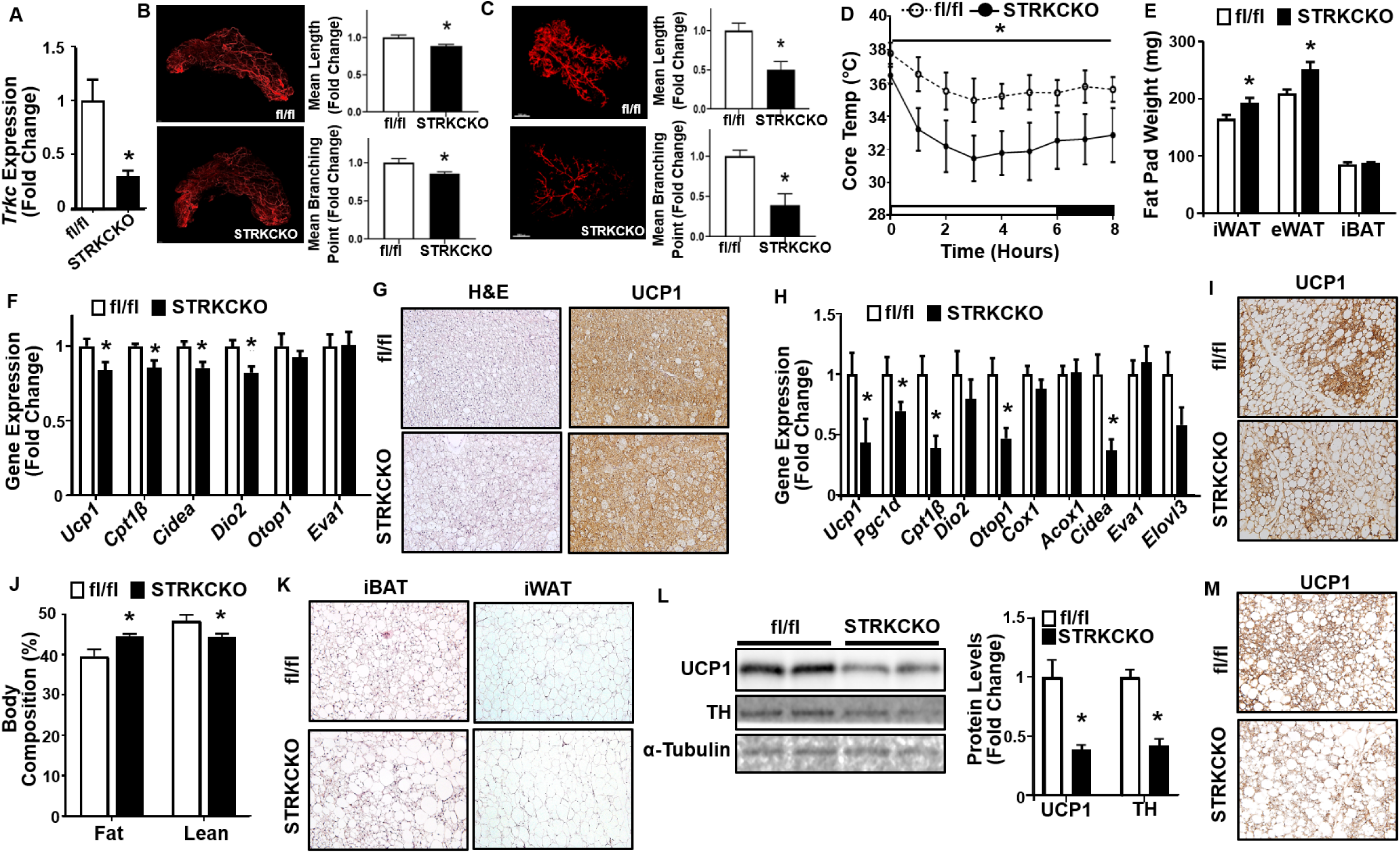
Mice with specific deletion of TRKC in sympathetic neurons exhibit impaired cold-induced thermogenesis and are prone to diet-induced obesity. (A) *Trkc* expression in sympathetic ganglia of fl/fl and STRKCKO mice (fl/fl n=4, STRKCKO n=5). (B)-C) Representative images of iWAT (B) and eWAT (C) TH-positive sympathetic nerve innervation (left panel) and quantitation of mean nerve fiber length and mean nerve fiber branching points normalized to total adipose tissue area (right panel) in fl/fl and STRKCKO mice after a 7-day cold challenge (n=3). (D)-(I) Core body temperature (D)(n=4), Fat pad mass (E)(n=4), *Ucp1* and other thermogenic gene expression in iBAT (F)(n=4), Representative H&E and UCP1 immunostaining in iBAT (G)(n=3), *Ucp1* and other thermogenic gene expression in iWAT (H) (fl/fl n=10, STRKCKO n=6), and representative UCP1 immunostaining images showing UCP1-positive beige adipocytes in iWAT (I)(n=3) of fl/fl and STRKCKO mice during a 7-day cold challenge. (J)-(M) Body composition (J)(n=5), representative H&E staining of iBAT and iWAT (K)(n=3), UCP1 and TH protein levels in iBAT (L) (fl/fl n=6, STRKCKO n=5), and representative UCP1 immunostaining in iBAT (M)(n=3) of fl/fl and STRKCKO mice fed a HFD. All data are expressed as mean ± SEM. *p<0.05.

During a chronic 7-day cold challenge, sympathetic neuron-specific deletion of TRKC resulted in significantly reduced mean sympathetic nerve fiber length and mean sympathetic nerve fiber branching point in iWAT and eWAT compared to that of fl/fl littermates (**Fig 7B-C**), indicating reduced sympathetic innervation in fat tissues. As a result, STRKCKO mice had significantly reduced body temperature in response to cold exposure (**Fig 7D**), but had higher fat mass in iWAT and eWAT depots (**Fig 7E**) compared to fl/fl mice. STRKCKO mice exhibited reduced *Ucp1* and other thermogenic gene expression (**Fig 7F**) and larger brown adipocytes with reduced UCP1 immunostaining in iBAT during cold exposure (**Fig 7G**). Further, STRKCKO mice had reduced *Ucp1* and other thermogenic gene expression (**Fig 7H**) and reduced UCP1-positive beige adipocytes in iWAT in response to the 7-day cold challenge (**Fig 7I**). These data indicate that sympathetic TRKC is important in regulating sympathetic innervation in fat tissue, and in cold-induced brown and beige adipocyte thermogenesis.

We then put STRKCKO mice on HFD at ambient room temperature. Body composition analysis by Minispec NMR revealed increased fat mass with decreased lean mass in STRKCKO mice, albeit there was no change of body weight (**Suppl. Fig 16A-C**). Similar to TRKC+/- mice, STRKCKO mice exhibited no change in food intake (**Suppl. Fig 16D**). However, STRKCKO mice displayed decreased energy expenditure as shown by decreased oxygen consumption and heat production (**Suppl. Fig 16E-F**), with no change in locomotor activity (**Suppl. Fig 16G**). The reduced energy expenditure in STRKCKO mice was associated with significantly reduced UCP1 and TH protein levels in iBAT, as well as larger brown adipocytes with reduced UCP1 immunostaining in iBAT (**Suppl. Fig 16H-I**). STRKCKO mice also had reduced thermogenic gene expression and larger adipocytes in iWAT (**Suppl. Fig 16J-K**). Consistent with their increased adiposity, these mice showed glucose intolerance and insulin resistance in GTT and ITT tests, respectively (**Suppl. Fig 16L-M**).

We also performed HFD feeding in these animals under thermoneutrality (30°C) to better assess diet-induced thermogenesis ^4, 46, 47^. STRKCKO mice similarly displayed higher fat mass and lower lean mass compared to their control mice (**Fig 7J**) despite no change in body weight (data not shown). Histological analysis also revealed that STRKCKO mice had enlarged adipocyte size in both iBAT and iWAT (**Fig 7K**), indicating increased lipid accumulation. In consistence, STRKCKO mice had significantly reduced UCP1 and TH protein levels in iBAT with reduced UCP1 immunostaining and larger adipocytes (**Fig 7L-M**). With increased adiposity, STRKCKO mice exhibited glucose intolerance and insulin resistance in GTT and ITT tests, respectively (**Suppl. Fig 17A-B**). In sum, our data suggest that TRKC signaling is crucial in regulating cold- and diet-induced thermogenesis and whole body energy homeostasis.

## Discussion

The premise of this study was derived from several prior observations. For one, Xue et al discovered the developmentally-induced beige adipocytes that appear transiently, peaking at postnatal day 20 and then disappearing thereafter toward adulthood ^11^. However, the fundamental question regarding the mechanisms underlying the induction and disappearance of the developmental beige adipocytes remains unanswered. Moreover, several lines of evidence have demonstrated a key role of SNS in BAT/beige thermogenesis ^12, 13, 14, 15, 16^. Factors that promote sympathetic innervation into BAT and WAT in response to developmental environmental cues (e.g. cold) are understudied. Here we discovered a novel fat-derived neurotrophic factor NTF3 and its receptor TRKC as key regulators of SNS growth and innervation in adipose tissue.

As stated above, CNS-originated activation of SNS that drives the brown and beige fat thermogenesis has been a major focus of recent investigation ^20^. However, prior studies have also demonstrated peripherally-derived neurotrophic factors in SNS growth and activation. The neurotrophic factors or neurotrophins, including nerve growth factor (NGF), brain-derived neurotrophic factor (BDNF), neurotrophin-3 (NTF3) and neurotrophin-4/5 (NT4/5), are a family of closely related proteins that act as survival factors for sympathetic and sensory neurons, and control survival, development and function of neurons in both the CNS and peripheral nervous system (PNS) ^27, 28, 50^. In peripheral nerve systems including SNS, neurotrophins are usually synthesized at a considerable distance from neurons by non-neuronal cells (targets) that are contacted by axons of these neurons innervating the target tissue ^27, 28, 50^. Neurotrophins are then retrogradely transported from the target along the axons into the neuronal cell body, where they regulate neuron function. This mode of action continues during development and throughout the adult life in order to maintain the normal differentiation, growth and function of the neuron and thus maintaining sufficient PNS innervation in target tissues ^27, 28, 50^.

The role of BDNF in the regulation of energy balance has been extensively studied, which was largely contributable to its regulation of food intake through CNS mechanisms ^51, 52, 53, 54^. It was reported that resistance training and combined high intensity exercising and resistance training increased plasma NTF3 levels in overweight subjects ^55^; and administration of NTF3 in *db/db* mice reduced their blood glucose level ^56^. In addition, circulating NTF3 level was reported to be negatively correlated with total cholesterol and low-density lipoprotein (LDL)-cholesterol levels ^57^, indicating a role of NTF3 in metabolic regulation. However, whether and how NTF3 regulates energy balance has been largely unknown.

A recent study from Spiegelman’s group demonstrated that S100, a BAT-derived secretory protein, promotes SNS innervation into adipose tissue. However, it is not clear whether S100 exerts the neurotrophic effect on SNS by itself or needs to work with other neurotrophic factors ^21^. Interestingly, a newly identified hepatokine Tsukushi (TSK) has been shown to suppress brown fat thermogenesis via down-regulating sympathetic innervation ^58^. However, the exact mechanism underlying the inhibitory effect of TSK on SNS innervation is not entirely clear. Our study demonstrates that NTF3 is a novel fat-derived neurotrophic factor that promotes SNS innervation through its direct effect on neurite growth, resulting in enhanced brown/beige fat thermogenesis and energy expenditure.

Existing literature supports an important role of target-derived NTF3 in the regulation of normal SNS neuron function, growth and target tissue innervation. Homozygous NTF3 deficient mice have significantly depleted peripheral sympathetic neurons in SG ^23^, and reduced norepinephrine levels in the heart ^24^. In addition, in homozygous NTF3 knockout mice, sympathetic fibers fail to invade the pineal gland and the external ear postnatally; whereas in NTF3 heterozygous mice, sympathetic fibers invade the pineal gland, but fail to branch properly and form a ground plexus ^59^. On the other hand, cutaneous overexpression of NTF3 increases sympathetic neuron number in the SG and enhances hair follicle innervation ^60^. We demonstrate here that BAT and WAT also produce NTF3 that regulates SNS innervation and activation in BAT and WAT in response to the developmental and environmental cues.

In the current study, we have used pharmacological NTF3 injection and genetic models to delineate the physiological function of NTF3. Combination of our AdNTF3-Tg, AdNTF3-KI and NTF3 injection models help provide an overall picture of the physiological role of the fat-derived NTF3 in the regulation of energy metabolism. For one, characterization of our transgenic AdNTF3-Tg mice that have overexpression of NTF3 in fat tissue without a systemic increase of NTF3 protein in the circulation demonstrates that increasing NTF3 locally in fat tissue should have a sufficient and direct effect on SNS innervation, increasing beige cell formation and thereby promoting energy expenditure. One the other hand, AdNTF3-KI and NTF3 injection models that have increased NTF3 in circulation appear to show a stronger phenotype of energy metabolism that of the AdNTF3-Tg mice, suggesting that NTF3 may have other beneficial effects systemically, for example, through the CNS that secondarily drives the SNS innervation into adipose tissue. It is noteworthy that such systemic effect of NT3 may align well with the therapeutic approach that often deliver agents systemically.

It appears that NTF3/TRKC’s signal may directly stimulate sympathetic neuron axonal growth, which may contribute to its stimulatory effect on SNS innervation into fat tissue. Through RNAseq analysis, we have identified novels genes in several important pathways involved in neuronal axonal growth are regulated in an opposite direction in SG of AdNTF3-Tg and TRKC+/- mice, including the coordination of actin filaments and microtubules, the dynamic cytoskeletal polymers and “building blocks” that promote shape change and locomotion during the peripheral nerve axonal outgrowth and elongation; family of proteins with GTPase activities that regulate actin motility, polymerization and stabilization during axonal initiation and elongation; regulation of glucose metabolism that provides energy needed for neuronal activity and axonal growth; protein sorting and transporting machineries; and proteins involved in axonal transport and translation of mRNAs ^34, 35^. These genes and pathways may represent novel NTF3/TRKC downstream targets that warrant future investigation.

In summary, we have discovered that the fat-derived neurotrophic factor NTF3 and its receptor TRKC are key regulators of SNS growth and innervation in adipose tissue. Our data indicate that NTF3 expression might be responsible for the induction of developmentally-induced beige adipocytes via sympathetic innervation in adipose tissue. We also demonstrate that NTF3 promotes cold-induced thermogenesis, enhances systemic energy expenditure and prevents diet-induced obesity.

## Methods

### Mice

Mice with haploinsufficiency of TRKC (TRKC+/-) were purchased from the Jackson Laboratory (#002481, Bar Harbor, ME). Mice with sympathetic-specific deletion of TRKC (STRKCKO) were generated by crossing TRKC-floxed mouse model (fl/fl) where the TRKC kinase ATP-binding domain is flanked with loxP sites ^48^ (Jackson Laboratory #022364) with TH-Cre mice where Cre-recombinase expression is under the control of the sympathetic marker tyrosine hydroxylase (TH) promoter ^49^(Jackson Laboratory #008601) to achieve specific deletion of TRKC in sympathetic neurons.

Reporter mice that express enhanced yellow fluorescent protein (EYFP) under the control of TRKC promoter (TRKC-Cre::R26-stop-EYFP) were generated by crossing TRKC-Cre mice ^61^ (NIH-supported Mutant Mouse Resource and Research Centers (MMRRC), #000364-UCD) with mice expressing a loxP-flanked STOP sequence followed by EYFP gene inserted into the Gt(ROSA)26Sor locus ^62^(Jackson Laboratory #006148).

To generate mice with NTF3 overexpression in adipocytes (AdNTF3-Tg), a bacterial artificial chromosome (BAC) containing the mouse adiponectin gene was used to create the adiponectin promoter-NTF3 overexpressing construct. Full-length coding sequence of the mouse NTF3 gene with a SV40 polyA signal sequence was PCR-amplified and inserted into the ATG position on exon 2 of the adiponectin gene in BAC using homologous recombination. The DNA fragment containing 5kb adiponectin promoter and NTF3 coding sequence was subcloned into pCR-Blunt, excised and microinjected into pronuclei of fertilized embryos of C57BL/6J mice at our Georgia State University Transgenic and Gene Targeting Core (Atlanta, GA). Transgenic founders were identified by PCR amplification of a ∼900bp fragment from tail DNA.

To alternatively achieve specific overexpression of *Ntf3* in adipose tissue, we used a gene targeting strategy to insert the *Ntf3* transgene into the *Rosa26* locus, which is the most used locus for the site-specific integration of transgenes due to a high efficiency of homologous recombination ^25^. The murine *Ntf3* cDNA was cloned into a *Rosa26* locus targeting vector ^25^ purchased from Addgene (Cambridge, MA). Key features of the complete targeting construct as depicted in **Suppl. Fig. 7A** include the following: 1) a strong CAG promoter allowing ubiquitous expression of cDNA construct placed behind it; 2) a loxP-flanked transcriptional blocker (Stop) that contains the transcription-blocking sequence; 3) the coding sequence for *Ntf3* with woodchuck hepatitis virus posttranslational regulatory element (WPRE); and followed by 4) a flippase (*Flp*) recognition target (FRT)-flanked neomycin selecting cassette (Neo). With this design, the strong CAG promoter can only drive *Ntf3* expression when the transcriptional block is removed by a tissue-specific Cre, therefore achieving tissue-specific over-expression. The DNA construct was electroporated into ES cells for homologous recombination targeting the endogenous *Rosa26* locus. About 30% of ES cell colonies were identified as positive by Southern blotting showing a wild-type band at 4.3kb and a targeted band at 5.5kb (**Suppl. Fig. 7B**). Correctly targeted ES cells were injected into blastocysts by our Transgenic and Gene Targeting Core (Georgia State University, Atlanta, GA). The resulting chimeric founders were crossed with C57BL/6J mice to achieve germline transmission of the knock-in allele, which were further bred with Flp mice ^63^(Jackson Laboratory, Stock No. 009086) to delete the neomycin cassette. The resulting Rosa-NTF3^fl/+^ mice were crossed with Adiponectin-Cre mice to generate mice with adipocyte-specific NTF3 knock-in mice (Adiponectin-Cre::Rosa-NTF3^fl/+^, or AdNT3-KI) by deleting the loxP-flanked Stop sequence.

For NTF3 injection experiment, postnatal C57BL/6J mice at age of day 20 were intraperitoneally (I.P.) injected with recombinant human NTF3 (50μg/kg body weight, R&D Systems, Minneapolis, MN) daily for 10 days, while 6 weeks old C57BL/6J mice on HFD were I.P. injected with the same dose of recombinant human NTF3 (OriGene Technologies, Rockville, MD) every day for the first two weeks, then three times per week for 8 weeks. Similar NTF3 doses and frequencies have been reported in the literature in preventing experimental pneumococcal meningitis-induced neuron loss ^64^ and in modulating nociceptive threshold and the release of substance P from rat spinal cord ^65^.

For Fast Blue injection, TRKC-Cre::R26-stop-EYFP, AdNTF3-Tg, TRKC+/- mice and their littermate controls were anesthetized via isoflurane and a dorsal or ventral 2 cm incision was made to expose IBAT or iWAT. The retrograde tracer Fast Blue (FB) (2%; Polysciences, PA) was injected with a microsyringe into 5-10 separate loci (1 μl/locus) of each adipose tissue. Animals were given 10-14 days to recover and allow for retrograde Fast Blue transport.

All animal procedures were approved by the Institutional Animal Care and Use Committee of Georgia State University.

### Metabolic measurement

All mice were housed with a 12/12 h light–dark cycle in temperature- and humidity-controlled rooms with free access to water and food. TRKC+/-, STRKCKO, AdNTF3-Tg mice and their respective littermate controls were fed either low fat (LF) (Research Diets D12450B, 10% calorie from fat) or high fat (HF) (Research Diets D12492, 60% calorie from fat) diet for up to 20 weeks. Some experiments were conducted under thermoneutrality (30°C) to avoid any non-shivering thermogenesis that may be induced by mild cold stress in mice housed under ambient room temperature (20-22°C) as we recently reported ^45^, which may serve as a confounding factor in assessing diet-induced thermogenesis ^4, 46^. Various metabolic phenotypes were characterized as follows. **1)** Body weight were monitored weekly. **2)** Food intake, energy expenditure and activity levels were measured using PhenoMaster metabolic cage systems (TSE Systems, Chesterfield, MO). **3)** Body composition was analyzed using a Minispec NMR body composition analyzer (Bruker BioSpin Corporation; Billerica, MA). **4)** Insulin sensitivity was determined by glucose tolerance and insulin tolerance tests (GTT and ITT, respectively) as we previously described ^66^. At the end of dietary treatment, BAT and WAT tissues were collected for further analysis of brown fat/beige adipocyte thermogenic program including gene expression, protein expression and immunohistochemistry.

### Cold exposure

TRKC-Cre::R26-stop-EYFP, AdNT3-Tg, TRKC+/-, STRKCKO mice and their respective littermate controls were subjected to a chronic 7-day cold challenge (5°C). At the end of experiment, tissues were collected for further analysis of brown fat/beige adipocyte thermogenic program including gene expression, protein expression and immunohistochemistry. In some experiments, core temperature was measured by a temperature transponder implanted into the peritoneal cavity as we previously described ^33^. Briefly, 2 weeks prior to the start of cold exposure, animals were anesthetized and a small incision was made along the ventral midline to expose the peritoneal cavity. Sterile temperature transponders (IPTT-300, BioMedic Data Systems, Seaford, DE) were implanted into the peritoneal cavity, and the incision was closed with sterile dissolvable sutures and sterile wound clips. After recovery, animals were subjected to a chronic 7-day cold challenge (5°C). Core body temperature was measured every 1-2 hours at the designated days during the cold exposure.

### Quantitative RT-PCR

Total RNA from adipose tissues and SG were isolated using the Tri Reagent kit (Molecular Research Center, Cincinnati, OH) ^66^. The expression of genes of interest was measured by a one-step quantitative RT-PCR with TaqMan Universal PCR Master Mix reagents (ThermoFisher Scientific, Waltham, MA) using an Applied Biosystems QuantStudio 3 real-time PCR system (ThermoFisher Scientific) as we previously described ^66^. The mRNA quantitation was further normalized by the housekeeping gene cyclophilin. The sequences of the primer and probe pairs for UCP1 and cyclophilin are as follows. UCP1: Forward 5’-CACCTTCCCGCTGGACACT-3’; Reverse 5’-CCCTAGGACACCTTTATACCTAATGG-3’; Probe 5’-AGCCTGGCCTTCACCTTGGATCTGA-3’. Cyclophilin: Forward 5’-GGTGGAGAGCACCAAGACAGA-3’; Reverse 5’-GCCGGAGTCGACAATGATG-3’; Probe 5’-ATCCTTCAGTGGCTTGTCCCGGCT-3’. The TaqMan primers/probes for all the other genes were purchased from Applied Biosystems (ThermoFisher Scientific).

### RNAseq analysis

RNA-seq library preparation, sequencing and basic bioinformatics data analysis from SG were performed by BGI Americas (Cambridge, MA), which specializes in high-throughput RNA/transcriptome sequencing and bioinformatics data analysis ^67, 68^. Briefly, after total RNA extraction and digestion with DNase I, mRNA were enriched with the oligo(dT) magnetic beads, fragmented (about 200 bp), and used for cDNA synthesis with random hexamer-primer. The double-stranded cDNA was ligated with sequencing adaptors and PCR amplified. RNA-seq libraries were sequenced using Illumina HiSeq™ 2000 (SE50). For quality control, RNA and library preparation integrity were verified using Agilent 2100 BioAnalyzer system and ABI StepOnePlus Real-Time PCR System.

For bioinformatics analysis, raw reads were filtered to remove adaptor sequences and low-quality data. Filtered clean reads were mapped to reference sequences (University of California Santa Cruz Mouse Genome Browser mm9 Assembly) using SOAPaligner/SOAP2 ^69^. Reads Per kilobase per Million reads (RPKM) was calculated to represent the gene expression level, and was used for comparing differentially expressed genes (DEGs) among groups. To identify genes regulated in the opposite direction in SG from thoracic T1 to T4 levels of AdNTF3-Tg and TRKC+/- mice after 1-day cold challenge, the RNA-seq datasets were merged based on Entrez gene ID. Genes up- or down-regulated in opposite direction in AdNTF3-Tg and TRKC+/- mice were identified using a cutoff value of absolute log_2_ fold change (log2FC) ≥ 0.5.

### Immunoblotting and NTF3 level measurement

Protein expression in adipose tissue was measured by immunoblotting as we described ^66, 70, 71^. Fat tissues were homogenized in a modified radioimmunoprecipitation assay (RIPA) lysis buffer supplemented with 1% protease inhibitor mixture and 1% phosphatase inhibitor mixture (Sigma-Aldrich, St. Louis, MO). Tissue lysates were resolved by SDS-PAGE. Proteins on the gels were transferred to nitrocellulose membranes (Bio-Rad, Hercules, CA), which were then blocked, washed, and incubated with various primary antibodies, followed by Alexa Fluor 680-conjugated secondary antibodies (Life Science Techenologies). The blots were developed with a Li-COR Imager System (Li-COR Biosciences, Lincoln, NE). The following primary antibodies were used: UCP1 (1:500, abcam, ab23841), TH (1:500; EMD Millipore, Temecula, CA), and α-Tubulin (1:1000, Advanced BioChemicals, ABCENT4777).

Tissue and serum NTF3 content was measured by ELISA (Advanced BioChemicals, Lawrenceville, GA).

### Immunohistochemistry (IHC)

Fat tissue was fixed in 10% neutral formalin, embedded in paraffin, and cut into 5 µm sections, which were either processed for hematoxylin and eosin (H&E) staining or immuno-staining with a UCP1 antibody (1:150, abcam, ab10983) as we previously described ^16, 33^. For immunostaining, the secondary antibody was conjugated with horseradish peroxidase (HRP) from Vector Labs (Vectastain ABC HRP kit, Vector Labs, PK-6100) and was developed with the 3,3’-diaminobenzidine (DAB) HRP substrate kit (Vector Labs, SK-4100).

SG were sectioned and immunostained as we described previously ^33^. Briefly, SG were carefully harvested, and transferred to an 18% sucrose solution in 0.1 M PBS containing 0.1% sodium azide at 4°C. All ganglia were then sectioned longitudinally at 10-µm-thick sections using a cryostat. They were directly mounted onto slides (Superfrost Plus; VWR International, West Chester, PA) in four series with every fifth section on the same slide. Sections were then incubated with rabbit anti-Tyrosin hydroxylase (TH, 1:1000; Millipore, MA) or rabbit anti-cFos (1:400; Millipore, MA) antibodies overnight and then incubated with donkey anti-rabbit Cy3 (1:400; Jackson Immunoresearch, West Grove, PA) antibodies for 2 h. Sections were mounted onto slides and cover slipped using ProLong Gold Antifade Reagent (Life Technologies, Grand Island, NY). Images were captured using an Olympus DP73 photomicroscope and CellSens software (Olympus, Waltham, MA). The captured images were evaluated with the aid of Olympus CellSens and the Adobe Photoshop (Adobe Systems, San Jose, CA) software. After two images were overlaid, exhaustive counts of FB- and TH (or cFos)-single neurons as well as FB with TH (or cFos)-colocalized neurons were performed by use of the manual tag feature of the Adobe Photoshop software in every fifth section of the ganglia to eliminate the likelihood of counting the same neuron twice. The neurons were considered positively labeled based on the fluorescent intensity, cell size, and shape. Percentage of positively single- and double-labeled neurons in the ganglia were averaged across each examined region from all mice.

### Assessment of sympathetic nerve innervation with adipose tissue whole mount clearing and immunostaining

A whole mount adipose tissue clearing with the Adipo-Clear approach ^22^ was conducted to allow immunostaining, followed by three dimensional visualization and quantitation of sympathetic nerve innervation by inverted confocal fluorescence microscopy. Briefly, adipose tissue was processed with a series of steps including dehydration, delipidation, and permeabilization, which was further stained with primary anti-TH antibodies (Millipore, Rabbit AB152) and secondary antibodies (Cy™3 AffiniPure Donkey Anti-Rabbit IgG (H+L), Jackson ImmunoResearch, 711-165-152). Tissues were further cleared with dibenzyl ether (DBE) and imaged with Zeiss 710 NLO Laser Scanning Confocal Microscope (Optical Microscopy Core, Georgia Institute of Technology, Atlanta, GA). Three-dimentional image reconstruction, sympathetic nerve fiber tracing and nerve density quantification including nerve branching/intersection points and average neurite length, were analyzed using the Imaris Image Analysis Software, and volume of each WAT was obtained using Imaris Surface tool to correct the size difference of each sample. (Oxford Instruments, imaris.oxinst.com).

### Sympathetic neuronal culture and neurite growth measurement

SG were dissected, digested with collagenase I followed by Trypsin digestion. Dispersed neurons were cultured in complete neurobasal media (Fisher 10888022) with 1X B-27 (Fisher 17504044) and 200µM L-glutamine (Fisher 25030149) in poly-D-lysine and Laminin coated dishes. Cells were cultured for 48 hours with or without saline or NTF3 (100ng/ml, R&D Systems, Minneapolis, MN) treatment, replated and further cultured for additional 3-6 hours with or without saline or NTF3 (100ng/ml) treatment. Similar NTF3 dose has been used in neuronal cultures previously ^72^. Cells were then immunostained with antibodies against neurite growth marker βIII-tubulin ^73^. Neurite growth was then quantitated by Neuron J ^74^. Between 60-160 neurons were analyzed in each group, and neurite number and neurite length were normalized by number of neurons in each group when doing comparisons between groups.

### Norepinephrine turnover measurement

Basal norepinephrine (NE) level and NE turnover rate (NETO) in adipose tissues were measured as we previously described ^33^. Briefly, mice were handled daily for one week prior to the start of the NETO experiment to adapt them to handling and reduce stress-induced NE release. On the day that NETO was measured, mice were subjected to cold exposure at 5°C for 16 h and NETO was measured during the last 4 h of the experiment. mice were injected intraperitoneally with α-methyl-p-tyrosine (250 mg/kg α-MPT; Sigma-Aldrich), an active competitive inhibitor for TH, which is the rate-limiting enzyme for NE production, thus preventing the synthesis of catecholamine. A supplemental dose of α-MPT (125 mg/kg) was given 2 h after the initial dose to ensure the inhibition of catecholamine synthesis. Two hours after the second α-MPT injection, mice were euthanized and adipose tissues were quickly harvested, weighed, frozen in liquid nitrogen, and stored at -80°C until NE extraction.

To obtain baseline NE values, one-half of hamsters from each group were euthanized without receiving α-MPT injections 4 h before the conclusion of the study. The adipose tissues were processed and extracted for NE with dihydroxybenzylamine (Sigma-Aldrich) as an internal control for extraction efficiency. NE content and NETO were measured as described previously ^75^. NETO was calculated using the following formula: k=(lg[NE]_0_ − lg[NE]_4_)/(0.434 × 4) and K = k[NE]_0_, where k is the constant rate of NE efflux, [NE]_0_ is the initial NE concentration, [NE]_4_ is the final NE concentration, and K = NETO.

### Statistical analysis

All data are expressed as mean ± SE. Differences between groups were analyzed for statistical significance by t test, one-way ANOVA and ANOVA with repeated measures followed by Bonferroni’s and Holm-Sidak’s post hoc tests as appropriate. Statistical significance is considered at *p <* 0.05.

## Acknowledgements

This work is supported by NIH grants R01DK107544, R01DK118106 and R01DK125081, and American Diabetes Association (ADA) grant 1-18-IBS-260to BX; NIH grants R01DK115740 and R01DK118106, and ADA grant 1-18-19 IBS-348 to HS; NIH grant R01DK116496 and ADA grant 1-18-IBS-346 to LY.

## Author contribution

XC performed experiments of AdNTF3-Tg mouse generation and characterization, TRKC+/- mice characterization, NTF3 injection in adult mice fed a high fat diet, all sympathetic innervation with Adipo-clear and imaging, sympathetic neuron culture and neurite growth, generation and characterization of STRKCKO mice, and performed the data analysis of the experiments; JJ performed experiments of AdNTF3-KI mouse generation and characterization, and performed the data analysis of the experiments; RW performed experiments with NT3 injection of postnatal P20 mice and characterization of TRKC+/- mice, and performed the data analysis of the experiments; QC performed sympathetic neuron culture experiments and data analysis (along with XC), QC, FL, KL and SW assisted in various experiments; HDS performed bioinformatics analysis of the RNAseq data; GS and LY contributed to study design, technical inputs and review/edits on manuscript; HS and BZ conceived and designed the study and wrote the manuscript.

