## Supplemental Figures and Legends for "Adipose tissue-derived neurotrophic factor 3 regulates sympathetic innervation and thermogenesis in adipose tissue"

Running Title: NTF3 regulates sympathetic innervation in adipose tissue.

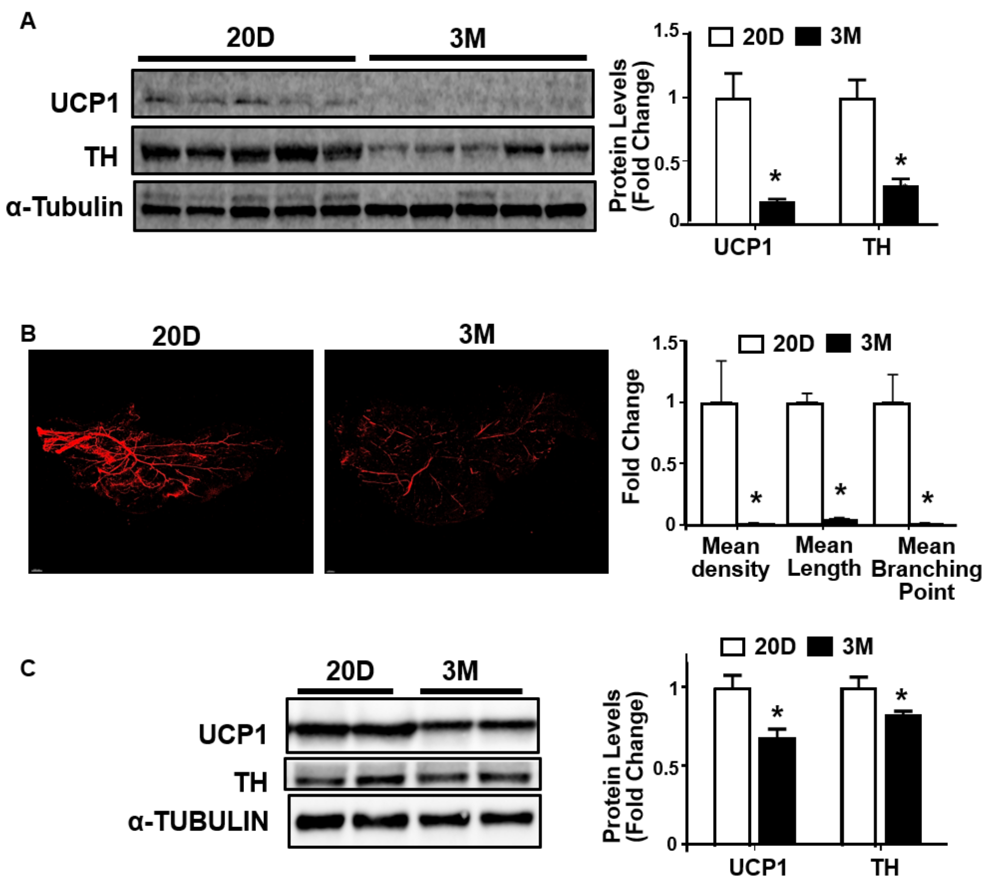


**Supplemental Figure 1.** Adipose-derived neurotrophic factor NTF3 correlates with UCP1 expression and sympathetic innervation in adipose tissue. (A) Immunoblotting (left panel) and quantitation (right panel) of UCP1 and TH protein in eWAT of 20 days old postnatal pups and 3-month-old adult mice (n=5). (B) Representative images of TH-positive sympathetic nerve innervation in eWAT (left) and quantitation of mean nerve fiber density, mean nerve fiber length, and mean branching points normalized to total adipose tissue area in eWAT (right panel) of 20 days old postnatal pups and 3-month-old adult mice (n=3). (C) Immunoblotting (left) and quantitation (right panel) of UCP1 and TH protein in iBAT of 20 days old postnatal pups and 3-month-old adult mice (20D n=4, 3M n=5). All data are expressed as mean ± SEM. *p<0.05.

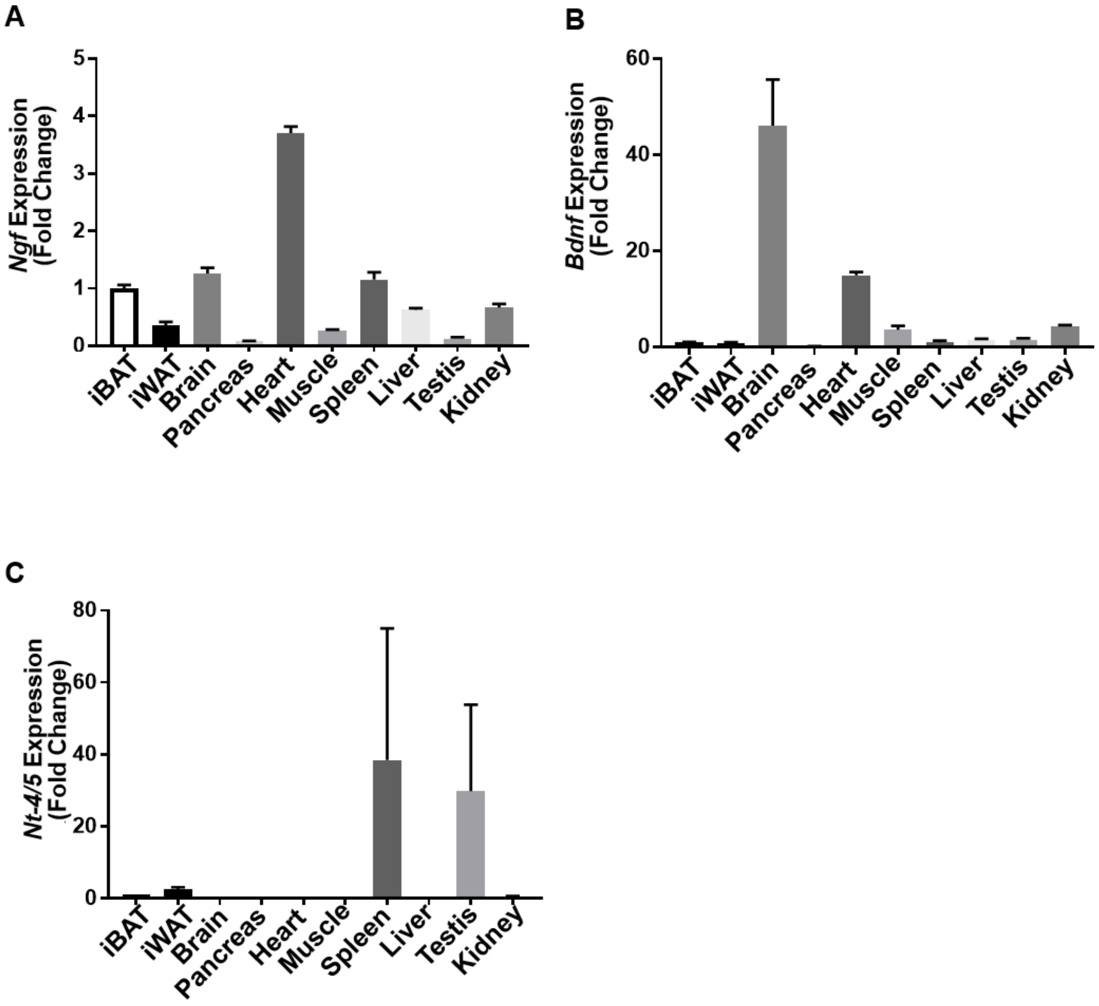


**Supplemental figure 2.** Tissue distribution of neurotrophic factors NGF (A), BDNF (B) and NT4/5 (C) (BAT n=7, WAT n=5, others n=4). All data are expressed as mean ± SEM.

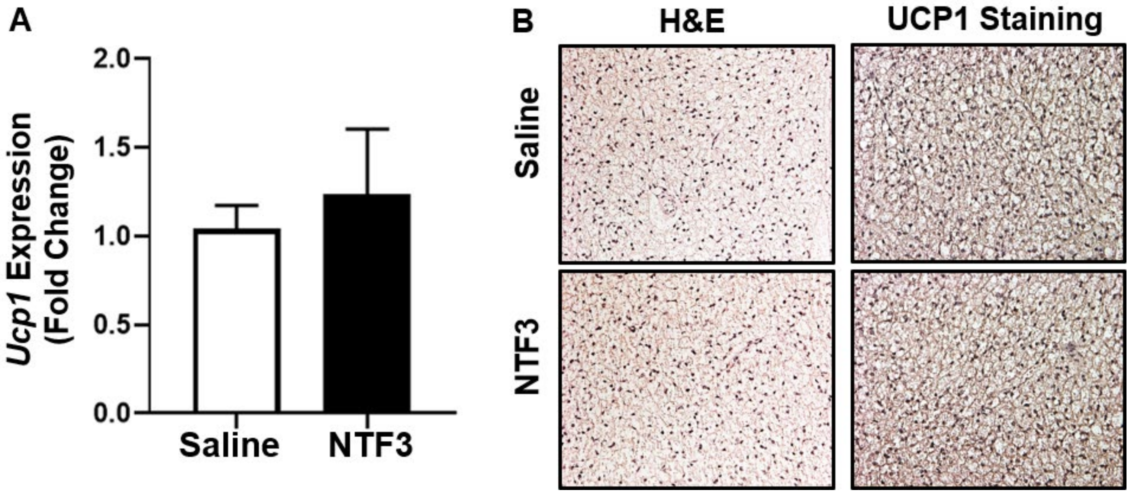


**Supplemental figure 3.** *Ucp1* expression (A)(n=5) and H&E and representative UCP1 immunostaining (B)(n=3) in iBAT of 20 days old postnatal pups with daily intraperitoneal (ip) NTF3 (50 μg/kg) injection for 10 days. All data are expressed as mean ± SEM.

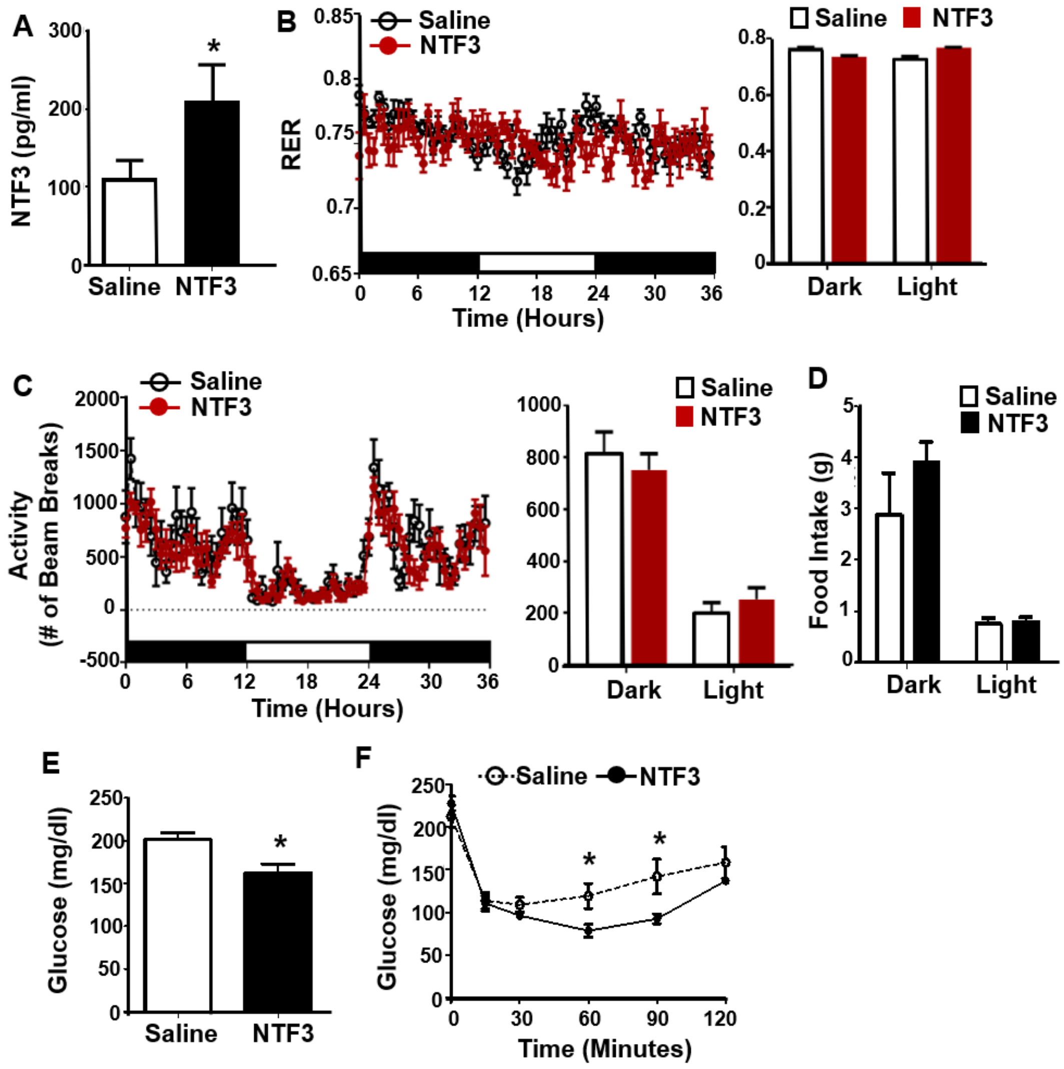


**Supplemental figure 4.** Metabolic characterization of NTF3-injected mice fed a HFD. Serum NTF3 levels (A) (saline n=8, NT3 n=6), Respiratory exchange rate (RER) (B)(n=8), Locomotor activity (C) (n=8), Food intake (D) (n=8), Fed glucose levels (E) (saline n=8, NTF3 n=6), and Insulin tolerance test (ITT) (F) (saline n=8, NTF3 n=6) in C57BL/6J mice injected with NTF3 with HFD feeding. All data are expressed as mean ± SEM. *p<0.05.

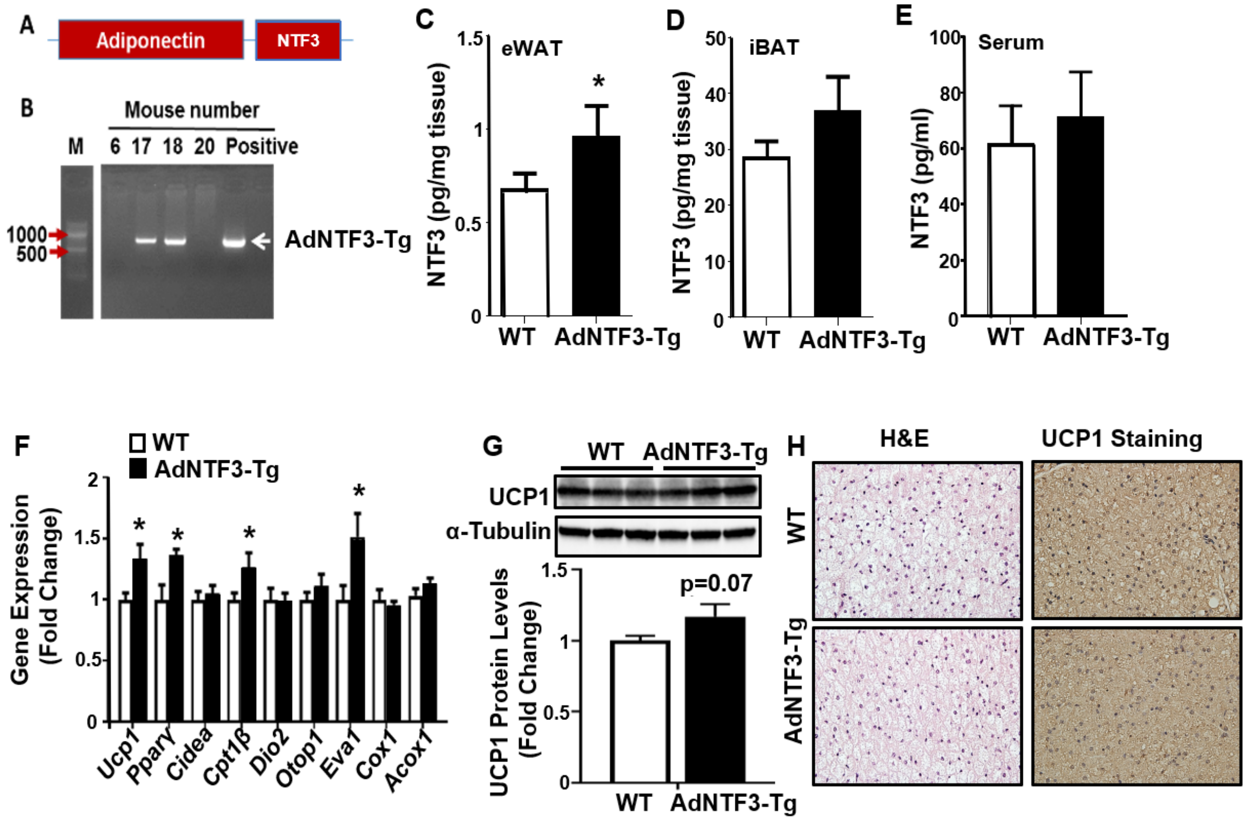


**Supplemental figure 5.** Generation and metabolic characterization of mice with adipocyte-specific overexpression of NTF3 (AdNTF3-Tg). (A) Schematic illustration of adiponectin-NTF3 overexpressing construct. (B) Genotyping identification of positive AdNTF3-Tg mice. (C)-(E) NTF3 protein levels in eWAT (C)(n=4), iBAT (D)(n=4) and serum (E) (n=4) of WT and AdNTF3-Tg mice as measured by ELISA. (F)-(H) *Ucp1* and other thermogenic gene expression (F)(n=5), UCP1 protein levels (G)(n=3) and representative H&E and UCP1 immunostaining in iBAT (H)(n=3) of WT and AdNTF3-Tg mice subjected to a 7-day cold exposure. All data are expressed as mean ± SEM. *p<0.05.

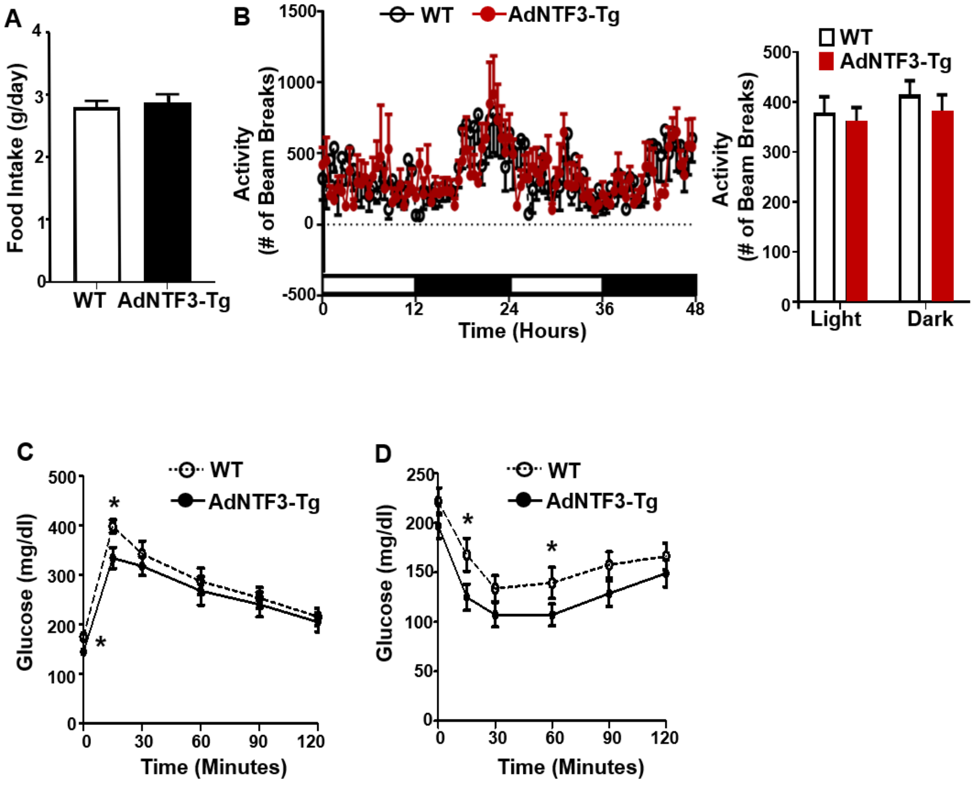


**Supplemental figure 6.** Metabolic characterization of WT and AdNTF3-Tg mice fed a HFD. Food intake (A)(n=4), Locomotor activity (B) (WT n=5, AdNTF3-Tg n=6), Glucose tolerance test (GTT) (C)(n=8) and ITT (D)(n=8) of WT and AdNT3-Tg mice fed a HFD. All data are expressed as mean ± SEM. *p<0.05.

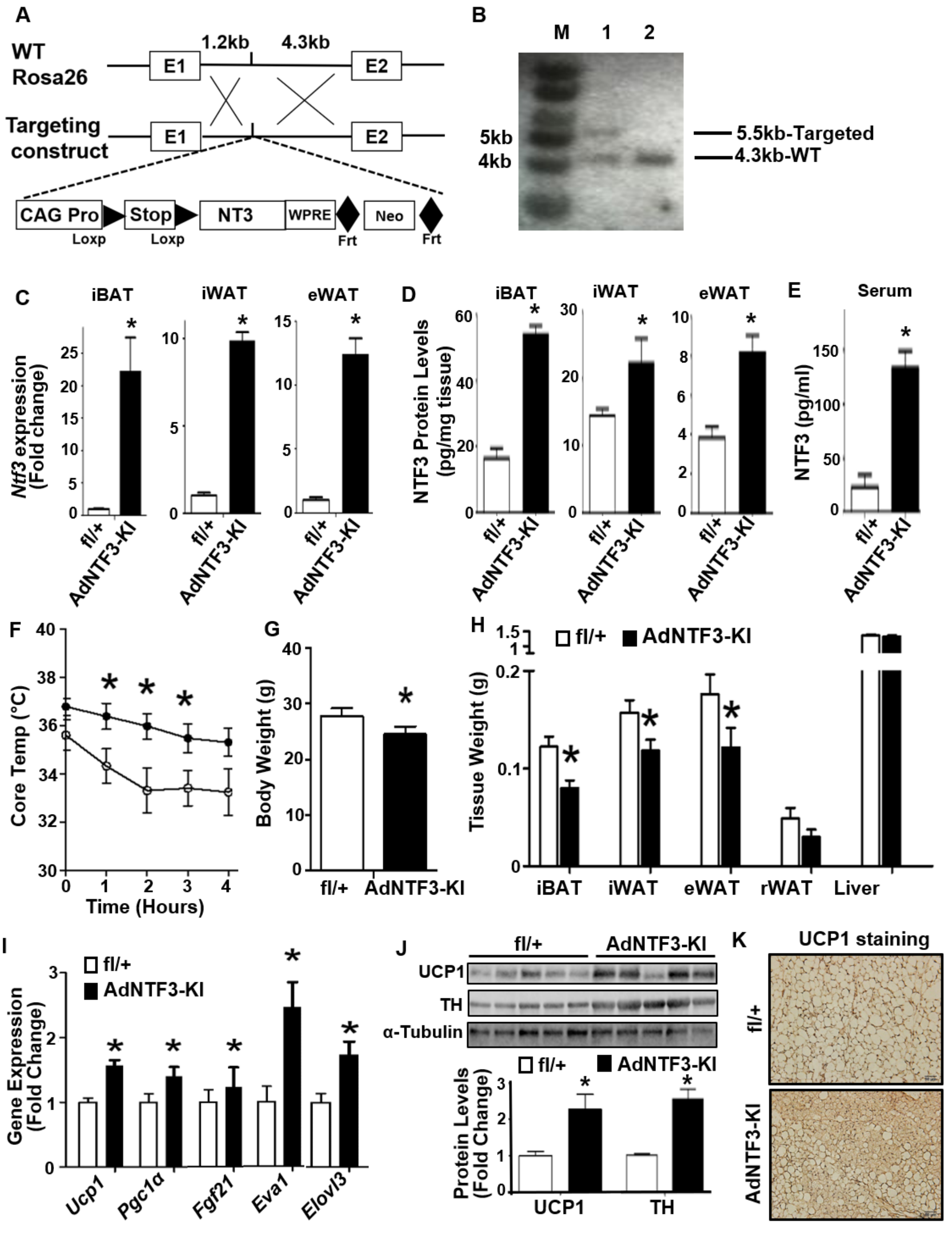
A
WT
Rosa26
1.2kb
4.3kb
E1
E2
Targeting
construct
CAG Pro
Stop
NT3
WPRE
Neo
Loxp
Loxp
Frt
Frt
B
M
1
2
5kb
4kb
5.5kb-Targeted
4.3kb-WT
C
iBAT
iWAT
eWAT
Ntf3 expression (Fold change)
fl/+
AdNTF3-KI
D
NTF3 Protein Levels (pg/mg tissue)
E
Serum
NTF3 (pg/ml)
F
Core Temp (°C)
Time (Hours)
G
Body Weight (g)
H
fl/+
AdNTF3-KI
Tissue Weight (g)
iBAT
iWAT
eWAT
rWAT
Liver
I
Gene Expression (Fold Change)
Ucp1
Pgc1α
Fgf21
Eva1
Elovl3
J
UCP1
TH
α-Tubulin
Protein Levels (Fold Change)
K
UCP1 staining
fl/+
AdNTF3-KI

**Supplemental figure 7.** Generation of AdNT3-KI mice with specific overexpression of *Ntf3* in adipocytes. (A) Schematic diagram of the gene targeting strategy to insert the *Ntf3* transgene into the *Rosa26* locus. CAG Pro: CAG promoter; Stop: transcriptional blocker; WPRE: woodchuck hepatitis virus posttranslational regulatory element; FRT: flippase (*Flp*) recognition target; Neo: neomycin selecting cassette. (B) Representative Southern blot for screening of homologous recombination of ES clones. (C)-(E) *Ntf3* mRNA (C) and protein levels (D) in fat tissue and serum (E) of 8 weeks old AdNTF3-KI and the control fl/+ mice (n=4). (F)-(K) Body temperature (F), Body weight (G), Fat pad mass (H), *Ucp1* and other thermogenic gene expression in iWAT (I), UCP1 and TH protein levels in iWAT (J) and representative UCP1 immunostaining image showing UCP1-positive beige adipocytes in iWAT (K) in 8 weeks old fl/+ and AdNTF3-KI mice in response to a 7-day cold challenge (n=5). All data are expressed as mean ± SEM. *p<0.05.

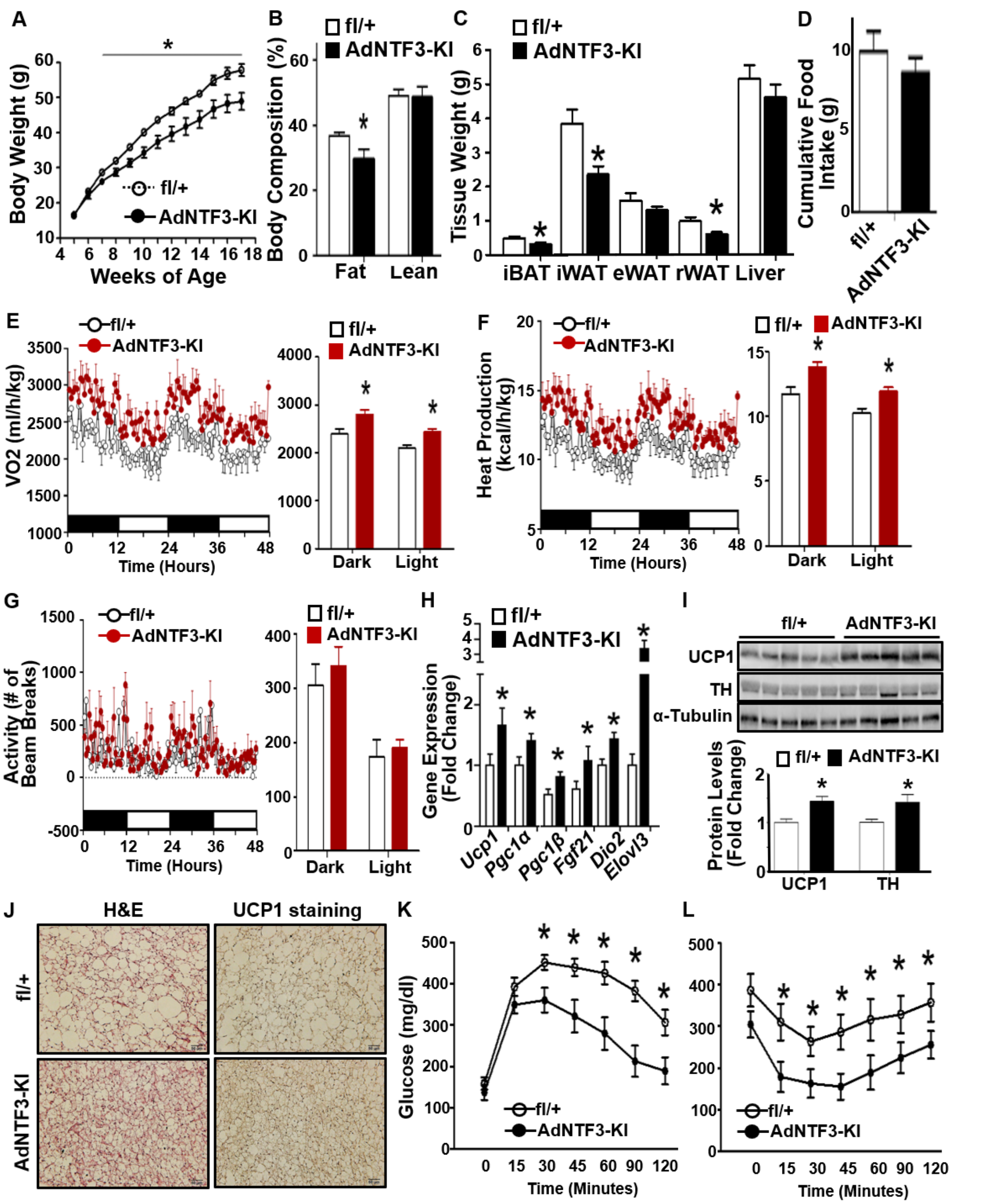


**Supplemental figure 8.** Metabolic characterization of fl/+ and AdNTF3-KI mice fed a HFD. Body weight (A), Body composition (B), Tissue weight (C), Cumulative food intake over 6 days (D), Oxygen consumption (E), Heat production (F), Locomotor activity (G), *Ucp1* and other gene expression in iBAT (H), UCP1 and TH protein levels in iBAT (I), Representative H&E and UCP1 immunostaining in iBAT (J), GTT (K) and ITT (L) in fl/+ and AdNTF3-KI mice fed a HFD (n=5). All data are expressed as mean ± SEM. *p<0.05.

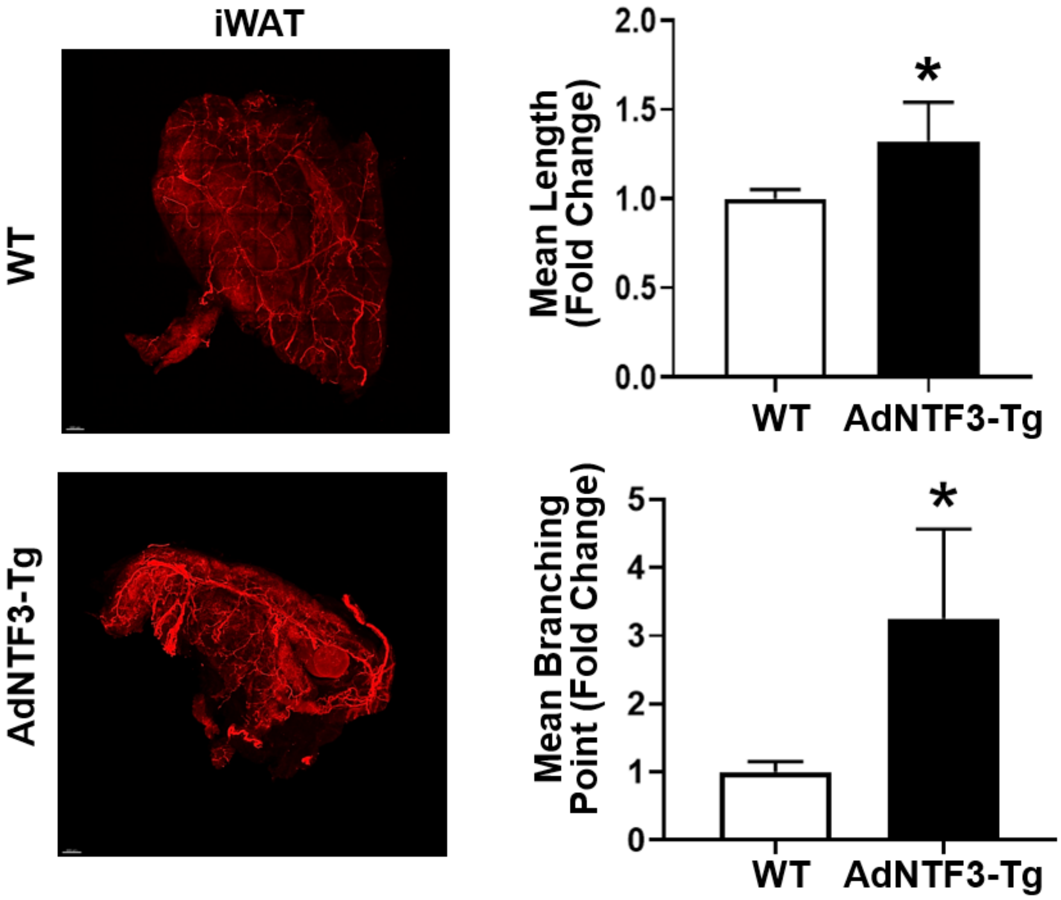


**Supplemental figure 9.** Representative images of iWAT TH-positive sympathetic nerve innervation (left panel) and quantitation of mean nerve fiber length and mean nerve fiber branching points normalized to total adipose tissue area (right panel) in 2-month-old WT and AdNT3-Tg mice housed at room temperature (n=3). All data are expressed as mean ± SEM. *p<0.05.

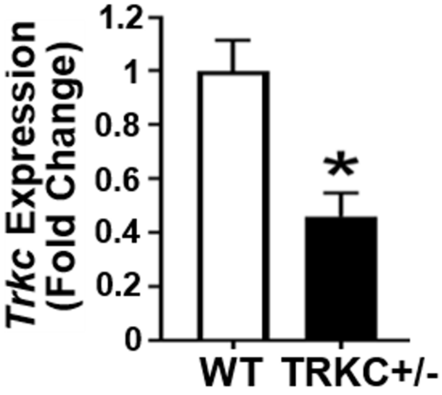


**Supplemental figure 10.** *Trkc* expression in sympathetic ganglia of WT and TRKC+/- mice (WT n=6, TRKC+/- n=4). All data are expressed as mean ± SEM. *p<0.05.

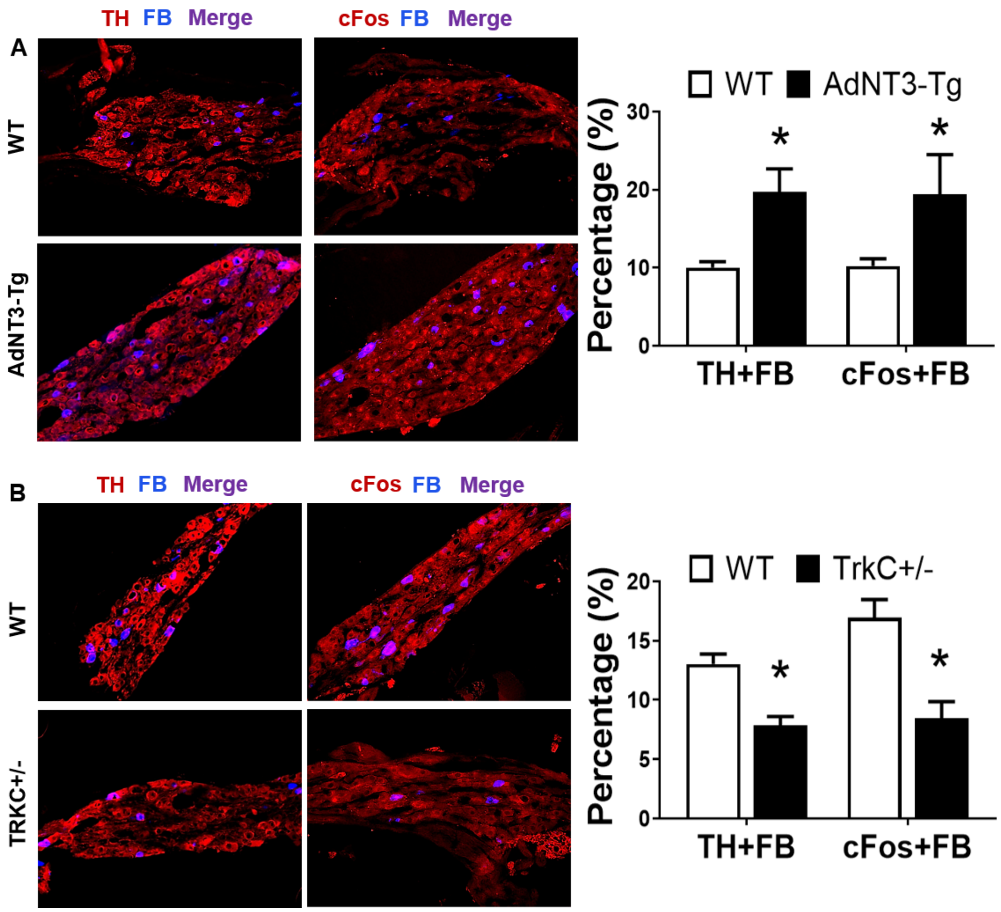


**Supplemental figure 11.** (A) Representative images (left panel) and quantitation (right panel) of TH/FB and cFos/FB labeling in sympathetic ganglia at lumbar L1 level in WT and AdNTF3-Tg mice after a 7-day cold challenge (n=4). (B) Representative images (left panel) and quantitation (right panel) of TH/FB and cFos/FB labeling in sympathetic ganglia at lumbar L2 level in WT and TRKC+/- mice after a 7-day cold challenge (n=4). All data are expressed as mean ± SEM. *p<0.05.

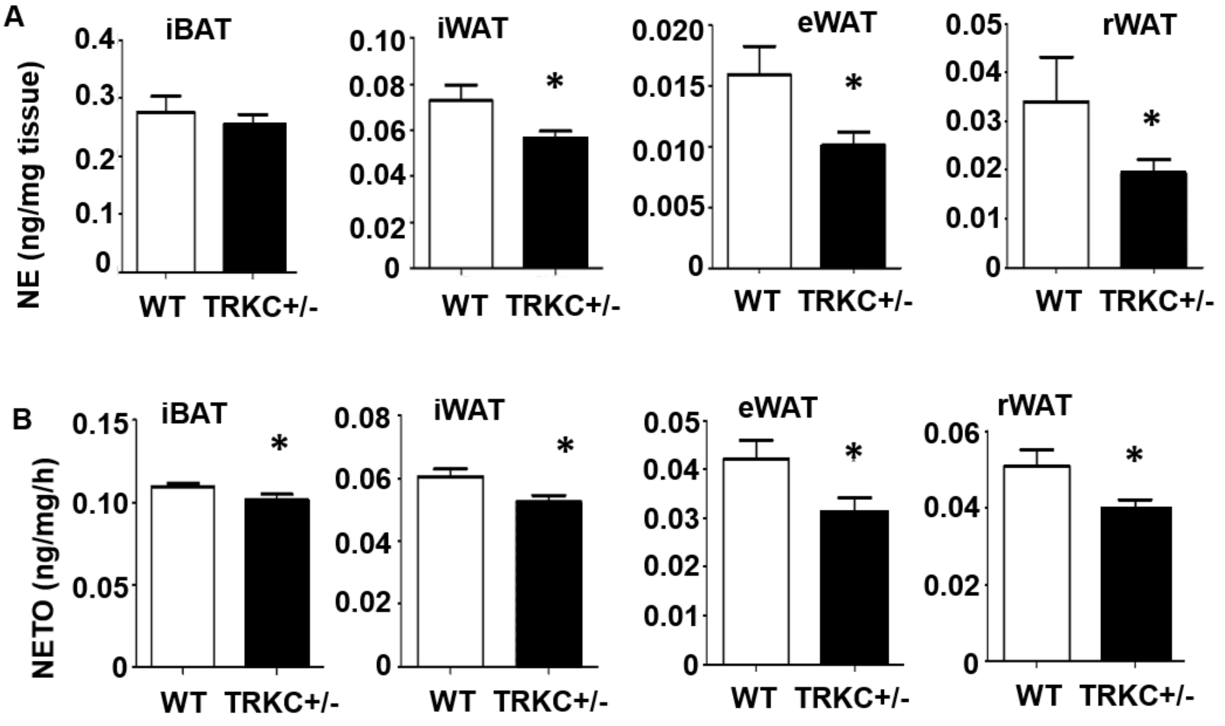


**Supplemental figure 12.** Basal norepinephrine (NE) content (A) and NE turnover (NETO) rate (B) in iBAT, iWAT, eWAT and rWAT of WT and TRKC+/- mice after a 16-hour cold challenge (n=7). All data are expressed as mean ± SEM. *p<0.05.

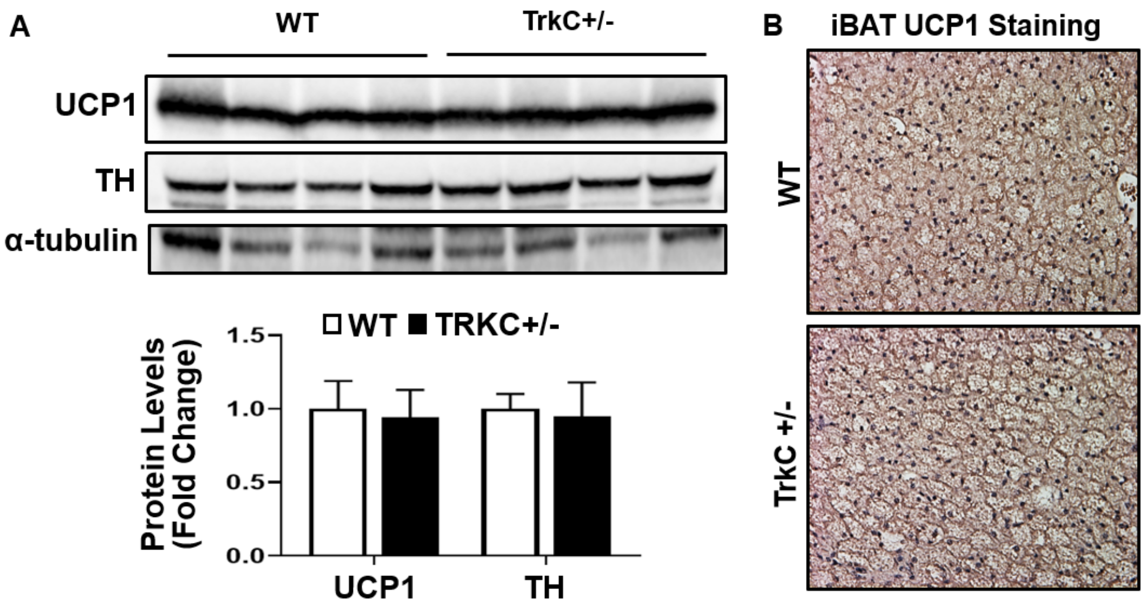


**Supplemental figure 13.** UCP1 and TH protein levels (A)(n=4) and representative UCP1 immunostaining in iBAT (B)(n=3) of 2-month-old WT and TRKC+/- mice subjected to a 7-day cold challenge. All data are expressed as mean ± SEM.

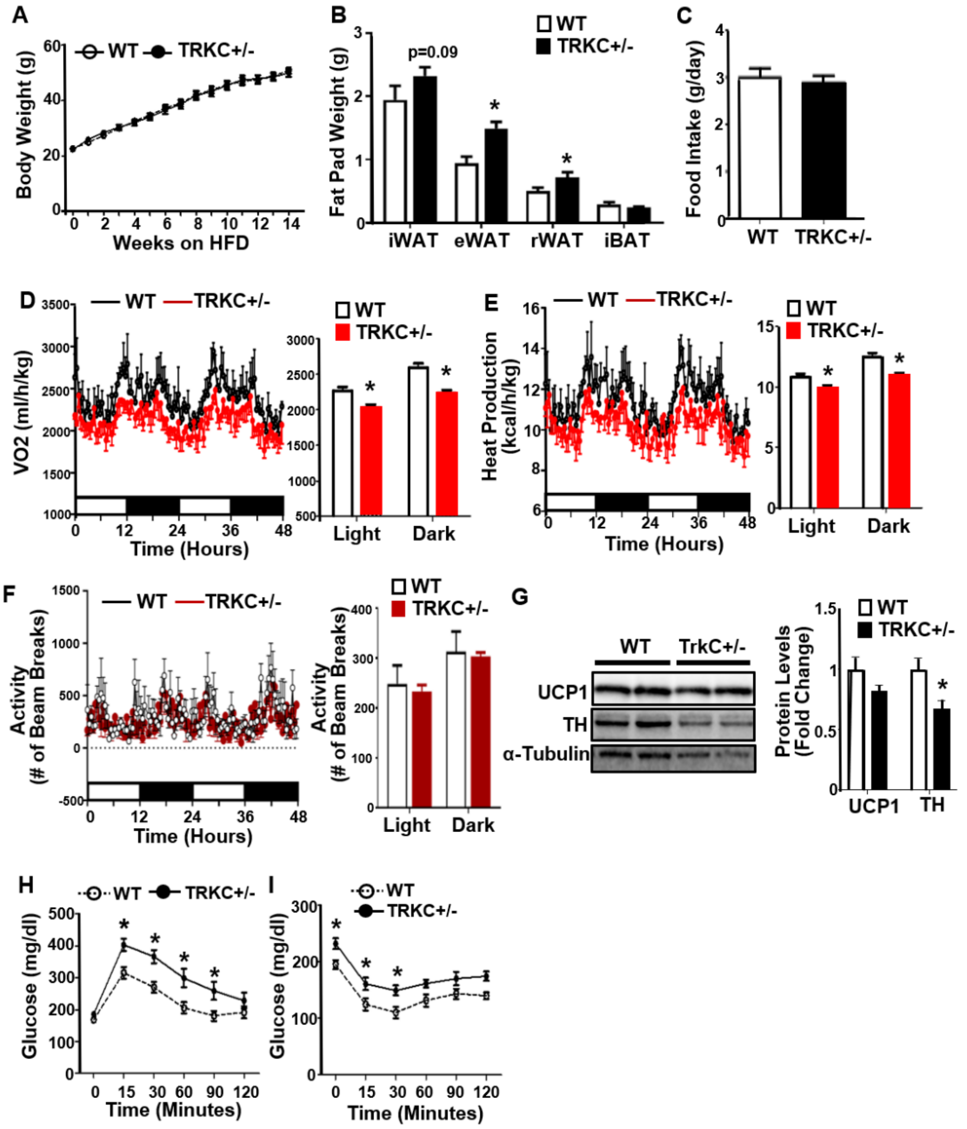


**Supplemental figure 14.** Metabolic characterization of WT and TRKC+/- mice fed a HFD at ambient room temperature (20-22°C). Body weight(A) (WT n=9, TRKC+/- n=7), Fat pad mass (B) (WT n=12, TRKC+/- n=9), Food intake(C) (WT n=8, TRKC+/- n=7), Oxygen consumption(D)(n=4), Heat production (E)(n=4), Locomotor activity (F)(n=4), UCP1 and TH protein levels in iBAT (G) (WT n=10, TRKC+/- n=4), GTT (H) (WT n=10, TRKC+/- n=7) and ITT (I) (WT n=8, TRKC+/- n=7) in WT and TRKC+/- mice fed a HFD at room temperature. All data are expressed as mean ± SEM. *p<0.05.

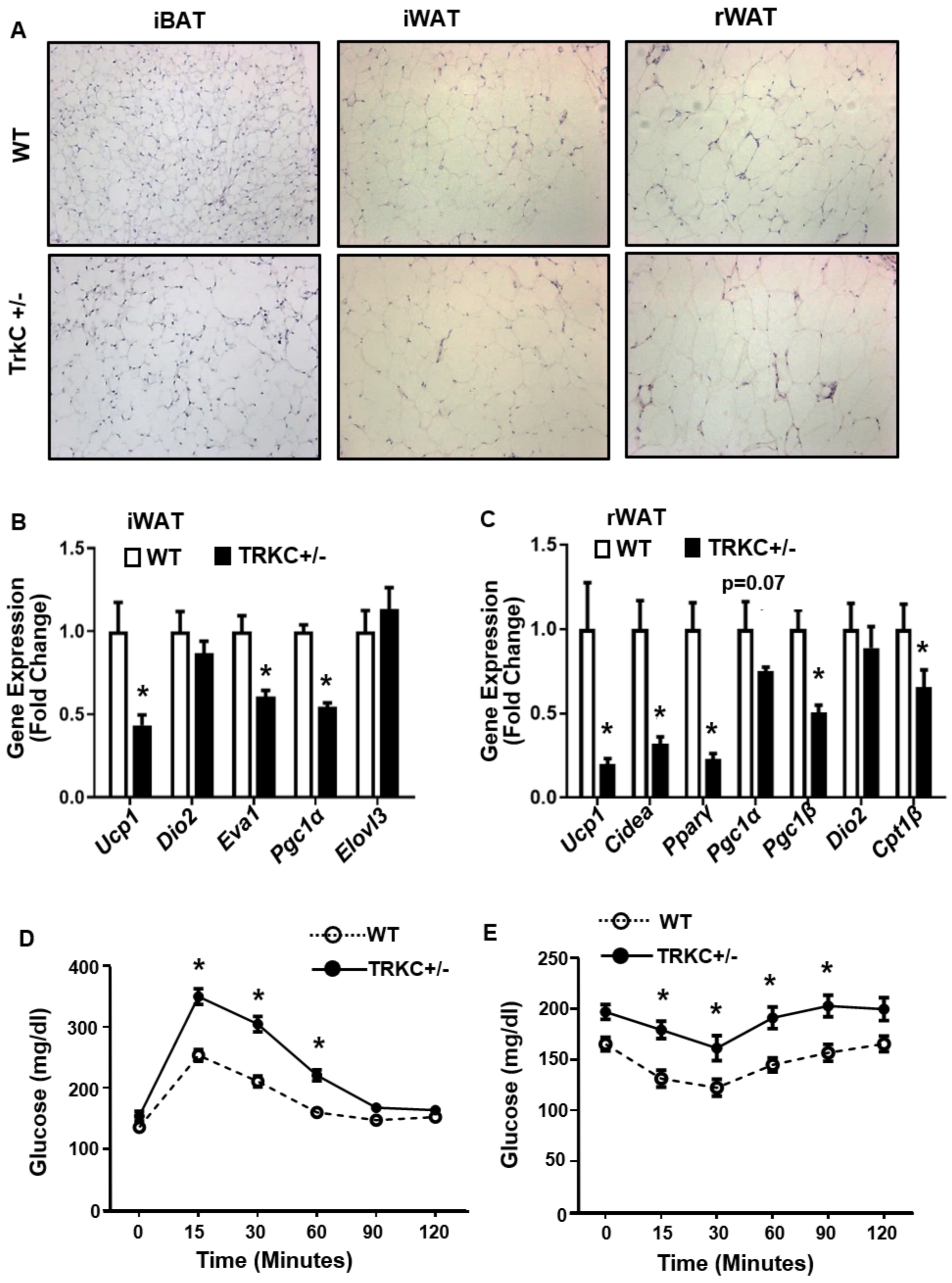


**Supplemental figure 15.** Metabolic characterization of WT and TRKC+/- mice fed a HFD at thermoneutrality (30°C). Representative H&E staining of iBAT, iWAT and rWAT (A)(n=3), *Ucp1* and other thermogenic gene expression in iWAT (B) (WT n=10, TRKC+/- n=9) and rWAT (C)(n=7), GTT (D) (WT n=14, TRKC+/- n=9) and ITT (E) (WT n=14, TRKC+/- n=9) in WT and TRKC+/- mice fed a HFD at thermoneutrality (30°C). All data are expressed as mean ± SEM. *p<0.05.

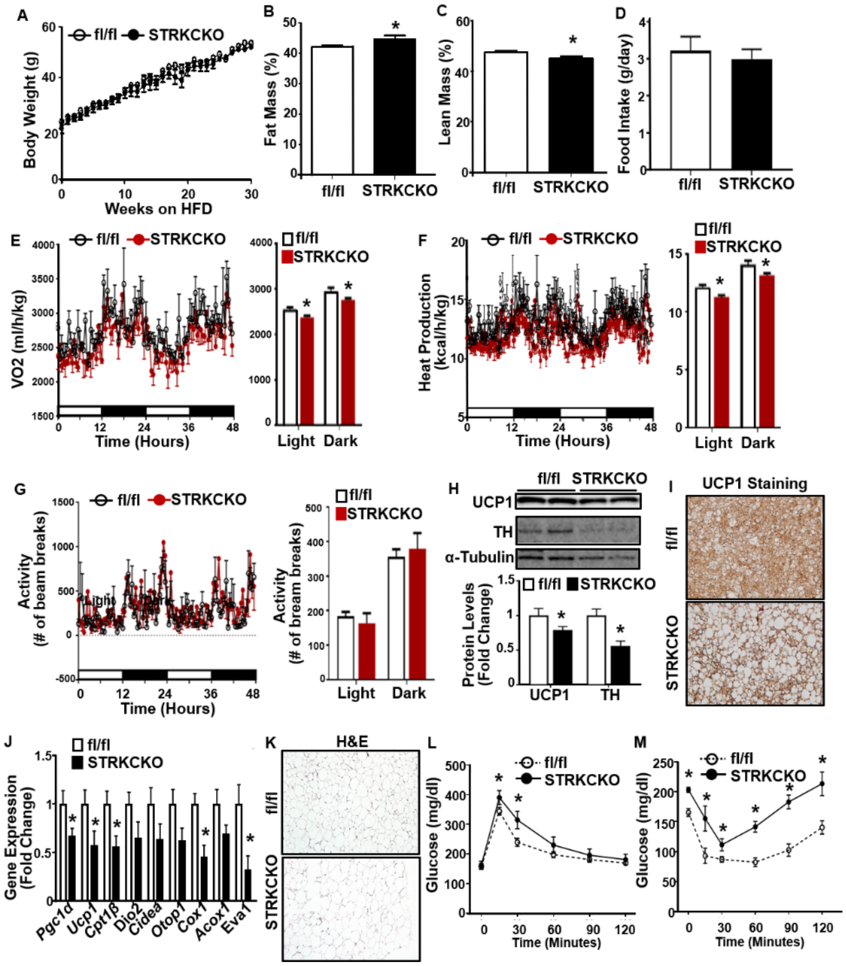

**Supplemental figure 16.** Metabolic characterization of fl/fl and STRKCKO mice fed a HFD at ambient room temperature (20-22°C). Body weight (A) (fl/fl n=8, STRKCKO n=6), Fat mass (B) (fl/fl n=8, STRKCKO n=6), Lean mass (C) (fl/fl n=8, STRKCKO n=6), Food intake(D)(n=6), Oxygen consumption(E)(n=6), Heat production (F)(n=6), Locomotor activity (G)(n=6), UCP1 and TH protein levels in iBAT (H)(fl/fl n=6, STRKCKO n=8), Representative UCP1 immunostaining in iBAT (I)(n=3), *Ucp1* and other thermogenic gene expression in iWAT (J) (fl/fl n=8, STRKCKO n=6), Representative H&E staining in iWAT (K)(n=3), GTT (L) (fl/fl n=8, STRKCKO n=6) and ITT (M) (fl/fl n=8, STRKCKO n=6) in fl/fl and STRKCKO mice fed a HFD at room temperature. All data are expressed as mean ± SEM. *p<0.05.

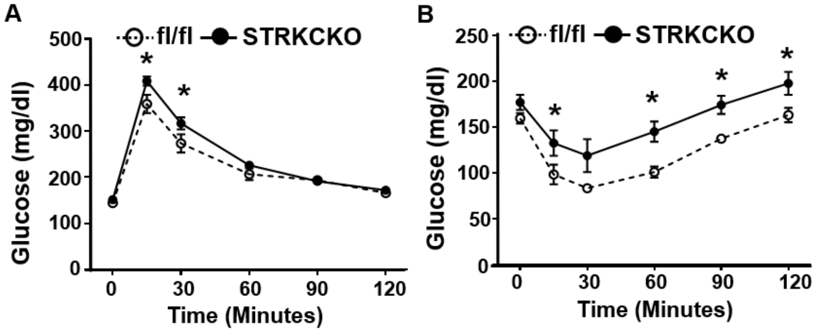


**Supplemental figure 17.** GTT (A) and ITT (B) in fl/fl and STRKCKO mice fed a HFD at thermoneutrality (30°C) (fl/fl n=5, STRKCKO n=6). All data are expressed as mean ± SEM. *p<0.05.
